# Missense variants in ACE2 are predicted to encourage and inhibit interaction with SARS-CoV-2 Spike and contribute to genetic risk in COVID-19

**DOI:** 10.1101/2020.05.03.074781

**Authors:** Stuart A. MacGowan, Geoffrey J. Barton

**Affiliations:** Division of Computational Biology, School of Life Sciences, University of Dundee, Dow Street, Dundee, DD1 5EH, Scotland, UK

**Author notes:** Correspondence should be addressed to Stuart A. MacGowan and Geoffrey J. Barton.

## Abstract

SARS-CoV-2 invades host cells via an endocytic pathway that begins with the interaction of the SARS-CoV-2 Spike glycoprotein (S-protein) and human Angiotensin-converting enzyme 2 (ACE2). Genetic variability in ACE2 may be one factor that mediates the broad-spectrum severity of SARS-CoV-2 infection and COVID-19 outcomes. We investigated the capacity of ACE2 variation to influence SARS-CoV-2 infection with a focus on predicting the effect of missense variants on the ACE2 SARS-CoV-2 S-protein interaction. We validated the mCSM-PPI2 variant effect prediction algorithm with 26 published ACE2 mutant SARS-CoV S-protein binding assays and found it performed well in this closely related system (True Positive Rate = 0.7, True Negative Rate = 1). Application of mCSM-PPI2 to ACE2 missense variants from the Genome Aggregation Consortium Database (gnomAD) identified three that are predicted to strongly inhibit or abolish the S-protein ACE2 interaction altogether (p.Glu37Lys, p.Gly352Val and p.Asp355Asn) and one that is predicted to promote the interaction (p.Gly326Glu). The S-protein ACE2 inhibitory variants are expected to confer a high degree of resistance to SARS-CoV-2 infection whilst the S-protein ACE2 affinity enhancing variant may lead to additional susceptibility and severity. We also performed *in silico* saturation mutagenesis of the S-protein ACE2 interface and identified a further 38 potential missense mutations that could strongly inhibit binding and one more that is likely to enhance binding (Thr27Arg). A conservative estimate places the prevalence of the strongly protective variants between 12-70 per 100,000 population but there is the possibility of higher prevalence in local populations or those underrepresented in gnomAD. The probable interplay between these ACE2 affinity variants and ACE2 expression polymorphisms is highlighted as well as gender differences in penetrance arising from ACE2’s situation on the X-chromosome. It is also described how our data can help power future genetic association studies of COVID-19 phenotypes and how the saturation mutant predictions can help design a mutant ACE2 with tailored S-protein affinity, which may be an improvement over a current recombinant ACE2 that is undergoing clinical trial.

**Key results:**

- 1 ACE2 gnomAD missense variant (p.Gly326Glu) and one unobserved missense mutation (Thr27Arg) are predicted to enhance ACE2 binding with SARS-CoV-2 Spike protein, which could result in increased susceptibility and severity of COVID-19
- 3 ACE2 missense variants in gnomAD plus another 38 unobserved missense mutations are predicted to inhibit Spike binding, these are expected to confer a high degree of resistance to infection
- The prevalence of the strongly protective variants is estimated between 12-70 per 100,000 population but higher prevalence may exist in local populations or those underrepresented in gnomAD
- A strategy to design a recombinant ACE2 with tailored affinity towards Spike and its potential therapeutic value is presented
- The predictions were extensively validated against published ACE2 mutant binding assays for SARS-CoV Spike protein

## 1 Introduction

The COVID-19 pandemic is one of the greatest global health challenges of modern times. Although the disease caused by the severe acute respiratory syndrome coronavirus 2 (SARS-CoV-2) is usually cleared following mild symptoms, it can progress to serious disease and death^1^. Besides the clear risks associated with age and comorbidities^2,3^, there could be a genetic component that predisposes some individuals to worse outcomes^4^. Identifying genetic factors of COVID-19 susceptibility has implications for clinical decision making and population specific epidemic dynamics. Genetic variation may constitute hidden risk factors and, in some cases, explain why otherwise healthy individuals in low-risk groups experience severe disease. The identification of specific genetic variants that influence the severity and progression of COVID-19 presents the opportunity for predictive diagnostics, early intervention and personalised treatments. The distribution of any genetic risk factors between populations could give rise to differences in population specific risk levels.

In the absence of GWAS hits for COVID-19 phenotypes, predictions of the effect that variation might have on the disease are essential. The emerging understanding of SARS-CoV-2 infection highlights proteins in which genetic variation could give rise to relevant phenotypic responses. Databases of human population variation provide a sample of variation in these proteins, to which we can apply state-of-the-art methods and workflows for predicting variant effects. A good candidate for variant analysis is a protein with a defined role in COVID-19 pathogenesis, with potentially functional variants reported in public databases and that has structural and functional data that can be used to interpret the potential effects of these variants.

Human angiotensin-converting enzyme 2 (ACE2) meets these criteria. ACE2 is the host cell receptor that SARS-CoV-2 exploits to infect human cells^1,5^. As this is the same receptor exploited by the SARS coronavirus (SARS-CoV) that caused the SARS outbreak in 2002, the detailed body of knowledge built around SARS-CoV infection is relevant to understanding SARS-CoV-2^1,5,6^. The spike glycoprotein (S) is the coronavirus entity that recognises and binds host ACE2. Both SARS coronavirus S-proteins are composed of an S1 domain that contains ACE2 recognition elements and an S2 domain that is responsible for membrane fusion^5^. The protease TMPRSS2 is the host factor that cleaves S into its S1 and S2 subdomains^5^, in SARS-CoV cleavage is thought to be promoted upon formation of the ACE2-S complex^7^. The S1 receptor binding domains (RBDs) from both SARS-CoV^8^ and SARS-CoV-2^9^ have been co-crystallised with human ACE2. The RBDs from both viruses are similar in overall architecture and both interface with roughly the same surface on ACE2. Differences are apparent in the so-called receptor binding motif (RBM) of S, which is the region of the RBD responsible for host range and specificity of coronaviruses^8–10^. The binding affinity of S and ACE2 is known explicitly to be correlated to the infectivity of SARS-CoV and is determined by the complementarity of the interfaces^8,10^.

In this paper we investigate the capacity of ACE2 variation to influence SARS-CoV-2 infection with a focus on predicting the effect of missense variants on the ACE2 SARS-CoV-2 S-protein interaction. The paper is organised as follows: In §2 we present an analysis of the effects of ACE2 missense variants on S-protein binding. We begin with an overview of missense variation at the ACE2-S interface from the Genome Aggregation Consortium Database (gnomAD^11^) and describe how these variants could modify binding with reference to structural and mutagenesis studies (§2.1). We then apply the mCSM-PPI2^12^ mutation effect predictor, which we have extensively validated against published ACE2 mutant SARS-CoV S-protein affinity assays^10^ (see Methods), to predict the effect of ACE2 variants from gnomAD (§2.2) and all possible missense mutations at ACE2-S binding residues (§2.3) on SARS-CoV-2 S-protein binding, and rationalise key results by a structural evaluation of the ACE2 variant models. In §3 we describe how other coding sequence variants from gnomAD^11^ might effect SARS-CoV-2 infection through mechanisms other than modulation of ACE2-S affinity and identify variants that may be of significance. In §4 we describe how this work could be applied to the development of an enhanced recombinant ACE2 therapy to treat COVID-19. Finally, in §5 we discuss the potential impact of ACE2 variants predicted to significantly affect SARS-CoV-2 infection in individuals and populations (§5.1), outline an application of our work to genetic association analyses (§5.2), address conflicting reports from similar studies and assess the confidence in our predictions (§5.3), and summarise our main results and conclusions (§5.4).

## 2 The effect of ACE2 missense variation on SARS-CoV-2 S binding

Figure 1 illustrates the distribution of gnomAD^11^ missense and nonsense variants in the sequence and structure^9^ of ACE2. In total, there are 338 ACE2 coding variants in gnomAD. The majority of these are missense variants (241, including 12 in splice regions) and synonymous variants (90, including 2 in splice regions). Synonymous variants do not alter the protein sequence and are therefore unlikely to have a functional impact. In general, the impact of missense variants is dependent on the structural and functional context of the mutated residue, as well as the physicochemical characteristics of the mutant residue. Missense variants are found in all the major functional domains of ACE2, including the peptidase and collectrin domains, the transmembrane region and other functionally annotated sites (Figure 1). Nonsense variants including stops and frameshifts are also reported in gnomAD for ACE2. These can have a drastic effect on protein structure and expression levels, but their effect is less dependent on the residue context than missense variants. However, there is linear positional dependence since truncations close to the C-terminus are more likely to be tolerated.

**Figure 1.**
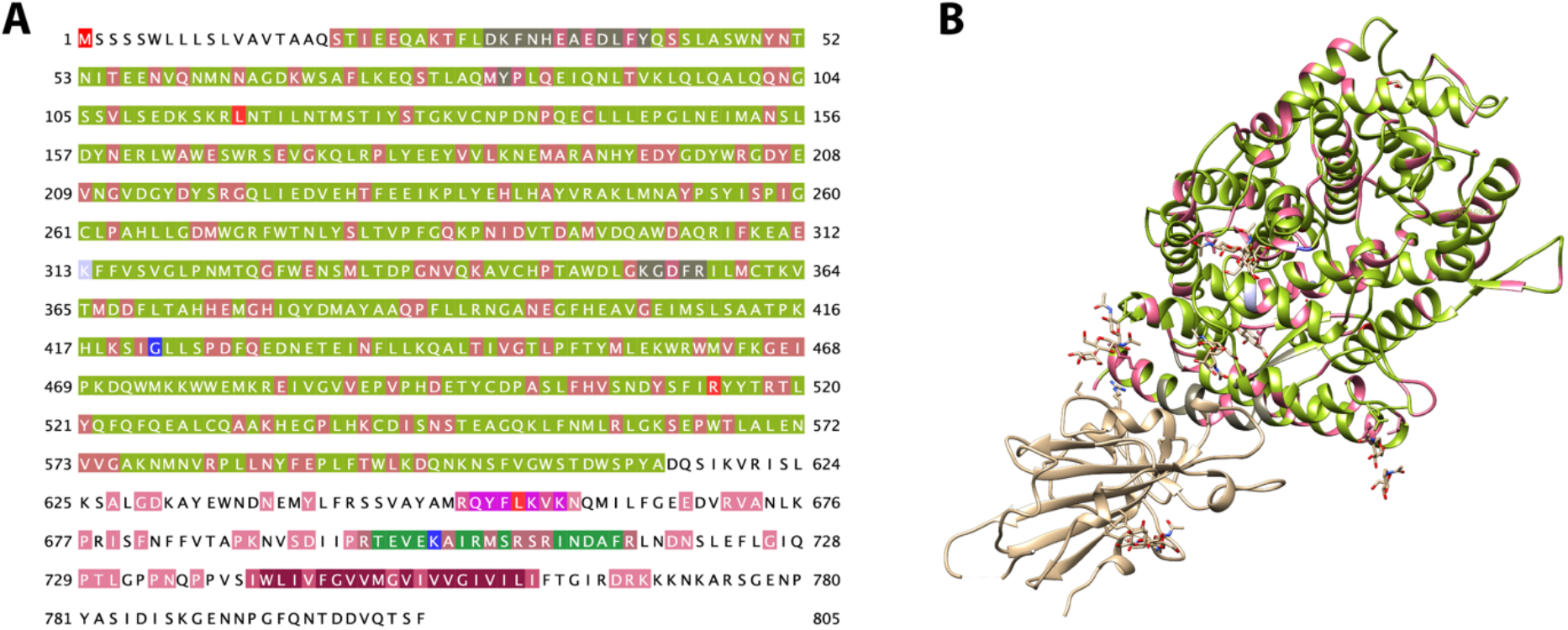
gnomAD missense variants in ACE2. A. gnomAD variants overlaid on the ACE2 sequence (UniProt Q9BYF1) and other sequence annotations. Highlights: missense variant (pink), nonsense variant (red), frameshift variant (blue), inframe deletion (light blue), coverage in PDB ID: 6vw1 (light green), SARS-CoV binding (dark grey, positions 30-41, 82-84 and 353-357), essential for ADAM17 processing (magenta, 652-659), essential for TMPRSS2 and TMPRSS11D processing (green, 697-716) and transmembrane region (dark red, positions 741-761). An expansion of this figure is given in Supplementary Figure 1. B. The ACE2-S complex from biological assembly 1 derived from PDB ID: 6vw1^9^. Sequence annotations retrieved from UniProt^13^ with Jalview^14^, variants were retrieved from gnomAD^11^ with VarAlign^15^. Figure generated with Jalview^14^ and UCSF Chimera^16^.

### 2.1 Missense variants and the ACE2-S interface

The simplest way that missense variation could impact SARS-CoV-2 infection would be by altering the ACE2-S interface. ACE2 missense variants located at residues that bind the S-protein are most likely to have such effects. Figure 2 illustrates the ACE2-S interface and highlights residues with reported missense variants in gnomAD whilst Table 1 lists these variants and their population distribution alongside key structural features of the site. Seven missense variants occur at residues we classified as directly interacting with SARS-CoV-2 S (see Methods). All these variants would be considered rare or extremely rare and are mostly private to a single gnomAD population. There are no homozygous females, but a few males are hemizygous as indicated. p.Ser19Pro is the most frequently observed ACE2-S interface variant in gnomAD. Mutations to or from Pro can have severe consequences for protein structure but as Ser19 is the first observed ACE2 residue in the structure it is difficult to predict the overall conformational consequence. This variant may be slightly deleterious to normal ACE2 function given the apparent gender imbalance in its prevalence, but the significance of this observation with respect to ACE2 activity towards SARS-CoV-2 infection is unclear. In the context of SARS-CoV-2 infection, the loss of the H-bond donor Ser, which interacts with a main chain carbonyl on S, could weaken the ACE2-S complex. The next most common missense variant at the interface is p.Glu37Lys. This variant occurs in an interface core binding site (see Methods), which have been shown to be less tolerant of mutation with respect to human pathogenic variants due to disruption of protein-protein interactions^17^. Charge inversion mutations have the capacity to be highly disruptive. Glu37 does interact with positively charged residues, but these are intraprotein interactions with ACE2 Arg393 and Lys353. Nevertheless, the excess positive charge in the variant may disfavour binding and the Lys will be unable to act as donor in the H-bond with SARS-CoV-2 S Tyr505 suggesting that this variant would be destabilising. Similar considerations hold for p.Glu35Lys. Glu35 engages with S via an H-bond to Gln493, which will be disrupted by the variant, but the effect of the charge inversion will be reduced compared to the previous variant because the neighbourhood of Glu35 includes both positive (Lys31) and negative (Glu38) charges. The site of p.Glu329Gly is close to the engineered S Arg439. Although it is not formally engaged in H-bonding or an ionic interaction by ARPEGGIO’s^18^ definitions, the residues are likely to interact electrostatically and has been described as an N…O bridge^9^. We would therefore expect ACE2 p.Glu329Gly to have reduced affinity with the chimeric receptor in PDB 6vw1. The wild-type Asn439 will likely be too short to interact with Glu329 and the loss of charge complementarity would probably lead to the same mutation having no effect with the native SARS-CoV-2 S (see Methods for further details).

**Figure 2.**
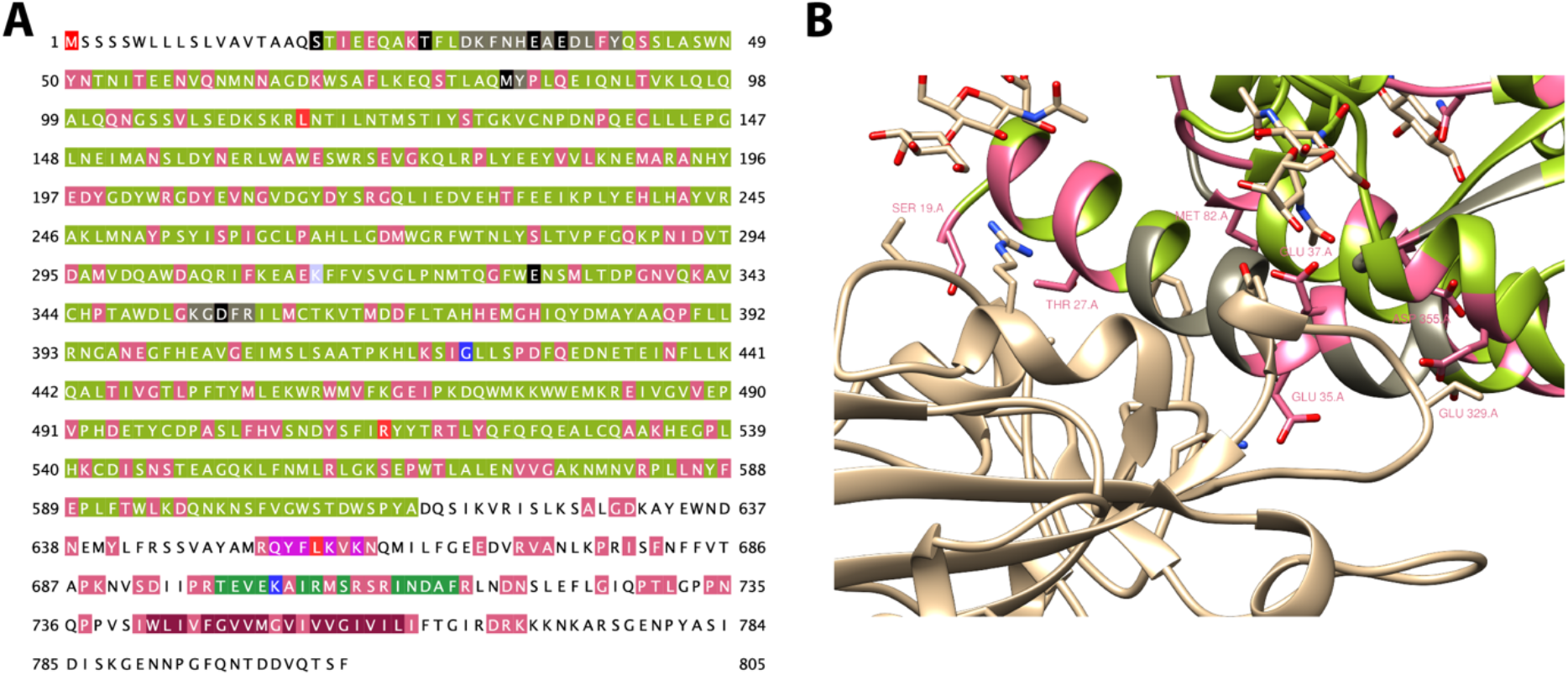
gnomAD missense variants at the ACE2-S interface. A. gnomAD variants overlaid on the ACE2 sequence (UniProt Q9BYF1) and other sequence annotations. Positions in ACE2 within 5 Å of S are highlighted with black backgrounds on the sequence (further annotations as in **Figure 1**). B. The corresponding residues are shown in the structure of the ACE2-S complex as resolved in PDB 6vw1^9^. Sequence annotations retrieved from UniProt^13^ with Jalview^14^, variants were retrieved from gnomAD^11^ with VarAlign^15^. Figure generated with Jalview^14^ and UCSF Chimera^16^.

**Table 1.** Frequency and structural features of ACE2 gnomAD^11^ missense variants at the ACE2 S-protein interface. Allele counts for males (AC Male) and the highest frequency population (AC Popn. Max) are provided alongside total allele counts (AC) to indicate whether a gender bias or population bias is present. Futher details on interface contacts are provided in Supplementary Table 1.

| ID | Mutation | Allele Count | AC Male | AC Popn. Max | Popn. Max<sup>a</sup> | Other Popn.<sup>a</sup> | Cons. | Contact Number<sup>b</sup> | Min. VdW Distance | Interface burial |
| --- | --- | --- | --- | --- | --- | --- | --- | --- | --- | --- |
| rs73635825 | p.Ser19Pro | 40 | 6 | 39 | AFR | OTH | 7 | 3 | 0.00 | Rim |
| rs781255386 | p.Thr27Ala | 2 | 0 | 2 | AMR |  | 8 | 5 | 0.40 | Rim |
| - | p.Glu35Lys | 3 | 1 | 2 | EAS | NFE | 7 | 1 | 0.00 | Rim |
| rs146676783 | p.Glu37Lys | 6 | 3 | 5 | FIN | AFR | 7 | 1 | 0.15 | Core |
| rs766996587 | p.Met82Ile | 2 | 0 | 2 | AFR |  | 9 | 1 | 0.08 | Rim |
| rs143936283 | p.Glu329Gly | 5 | 0 | 4 | NFE | OTH | 4 | 1 | 0.99 | Rim |
| - | p.Asp355Asn | 2 | 0 | 2 | NFE |  | 8 | 2 | 0.20 | Core |

Column legend – Popn. Max: *gnomAD* population with highest allele frequency: African/African-American (AFR), Latino/Admixed American (AMR), East Asian (EAS), Finnish (FIN), non-Finnish European (NFE) and Other/not-assigned (OTH). Other Popn.: other *gnomAD* populations in which the variant was also observed. Cons.: Physicochemical conservation of the mutation<sup>19</sup>, Contact Number: The number of S-protein residues the wild-type ACE2 residue interacts with. Interface burial: Whether the residue is situated at the interface core or rim.

The effects of ACE2 residue substitutions on coronavirus S-protein affinity have been directly observed^10^. Extensive ACE2 mutagenesis provided detailed insight into the ACE2-SARS-CoV S complex and can give some insight into the effects of ACE2 population variation in SARS-CoV-2 S binding. Table 2 lists gnomAD missense variants seen at the site of residues that were previously mutagenized^10^. Half of the ACE2 mutants listed resulted in inhibition of S-protein binding but only two mutants correspond exactly to the gnomAD variant and of these only p.Arg559Ser showed an effect on SARS-CoV S-protein binding. The other mutant also seen in gnomAD, p.His239Gln, had no reported effect on the interaction. The variant p.Lys68Glu shares physicochemical properties with the mutant Lys68Asp, which slightly inhibited interaction with SARS-CoV, and the loss of Pro389 that results from the variant p.Pro389His also had a slight reduction in binding from a Pro389Ala mutant. The only variant that is situated at an ACE2-S interacting residue and occurs at a site with a reported single-point mutant is p.Asp355Asn. The mutant Asp355Ala was reported to reduce ACE2 binding to SARS-CoV S-protein to about 20 % of the wild-type^10^. This indicates that the site is important for maintaining the interaction between the proteins.

**Table 2.**
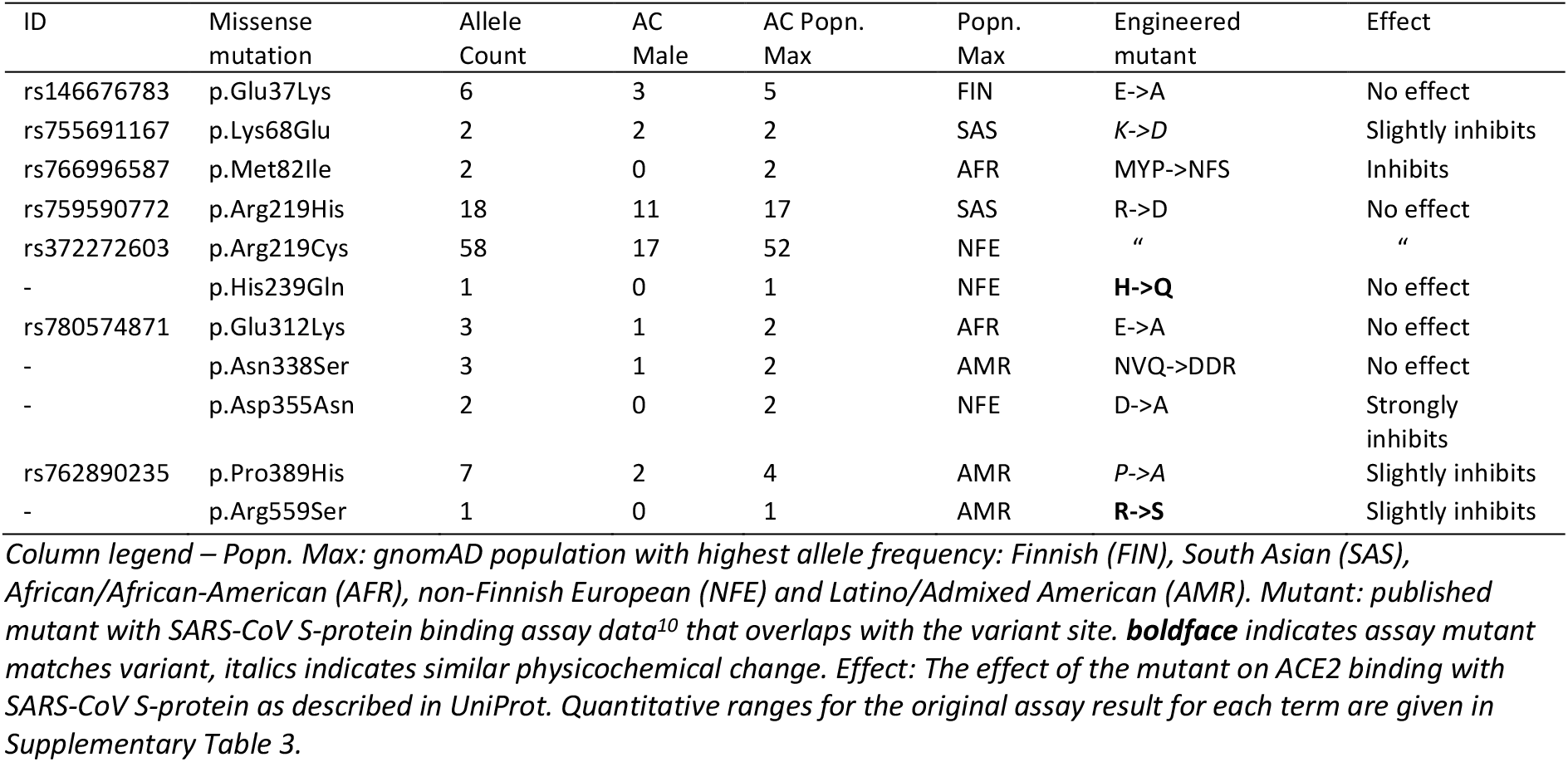
Frequency and site mutagenesis report for ACE2 gnomAD^11^ missense variants located at residues with previously reported ACE2 mutant SARS-CoV S binding assay^10^.

There are a few caveats to the preceding analyses. The first is that our structural consideration of variants located at the interface is qualitative. For the variants that we think would be destabilising, we are unable to say how destabilising they might be and would have difficulty even ranking them. The second is that we have not ascertained how far the assay results for SARS-CoV S are applicable to SARS-CoV-2 S. Structural comparison of the two complexes could help to address this second caveat since ACE2 sites that interact with S-protein sites that are conserved between SARS-CoV and SARS-CoV-2 are more likely to respond in the same way to a mutation than where SARS-CoV-2 has a different residue. A third flaw is that we have not considered all the reported missense variants. Although variants located at the interface are most likely to have an effect, residues in proximity to the interface, but not directly a part of it, could also have an effect. Second and third degree contacts can have an effect too and even longer range substitution induced conformational effects cannot be discounted completely (although it has been shown that SARS-CoV S binding is invariant to ACE2 catalytic conformational changes^8,10^). In §2.2 we address these caveats by employing mCSM-PPI2^12^ to predict changes in the ACE2-S binding energy as a result of the gnomAD variants. We integrate the experimental mutagenesis data by validating mCSM-PPI2 in this system and calibrating its predictions to the experimental observations.

### 2.2 gnomAD variants in ACE2 can significantly modify its affinity with SARS-CoV-2 S

Research into the impact of missense variants at protein-protein interaction interfaces has led to methods that provide quantitative predictions of the effect of mutations on interface characteristics^12^. These predictions are an invaluable addition to insights gained from inspection of the site and its neighbourhood in the 3D structure. They can also be used to screen large numbers of variants that would be prohibitively time-consuming to process through modelling and manual inspection or experimentally. In this section, we predict the effects of ACE2 missense variation on S-protein binding by the application of the mCSM-PPI2^12^ algorithm, which we extensively validated against experimental binding data for 26 ACE2 mutants in complex with the closely related SARS-CoV S (see Methods), and critically evaluate the key predictions by structural inspection.

Table 3 contains mCSM-PPI2 predictions for nine ACE2 gnomAD missense variants located at or close to the S-protein interface. On the basis of our mCSM-PPI2 calibration, predicted ΔΔG > +1.0 kcal/mol is a confident indication that a variant will enhance ACE2-S binding whilst ΔΔG < −1.0 kcal/mol indicates a mutation that strongly or totally abolishes the ACE2-S interaction (see Methods). One variant is predicted to enhance binding (p.Gly326Glu) whilst three variants are predicted to strongly reduce S-protein binding. The remaining five interface variants are predicted to reduce ΔG over the range −0.2 kcal/mol to −0.6 kcal/mol, but as they do not exceed the calibrated threshold, we consider these variants as of uncertain significance with respect to ACE2-S binding.

**Table 3.** mCSM-PPI2^12^ ΔΔG predictions and published mutagenesis assays^10^ for gnomAD^11^ missense variants at the ACE2 S-protein interface. See Supplementary Table 2 for mCSM-PPI2 predictions for all gnomAD variants mapped to PDB 6vw1.

| ID | Missense mutation | Distance to Interface | mCSM-PPI2 $\Delta\Delta G$ | Prediction | Mutagenesis |
| --- | --- | --- | --- | --- | --- |
| rs73635825 | p.Ser19Pro | 2.6 | -0.2 |  |  |
| rs781255386 | p.Thr27Ala | 3.7 | -0.6 |  |  |
| - | p.Glu35Lys | 2.9 | -0.5 |  |  |
| rs146676783 | **p.Glu37Lys** | **3.2** | **-1.2** | **Decreased affinity** | No effect (E->A) |
| rs766996587 | p.Met82Ile | 3.5 | -0.3 |  | Inhibits (MYP->NFS) |
| rs759579097 | **p.Gly326Glu** | **5.5** | **1.0** | **Enhanced affinity** |  |
| rs143936283 | p.Glu329Gly | 4.1 | -0.4 |  |  |
| rs370610075 | **p.Gly352Val** | **5.4** | **-1.1** | **Decreased affinity** |  |
| - | **p.Asp355Asn** | **3.5** | **-1.3** | **Decreased affinity** | Strongly inhibits (D->A) |

*Column legend – Distance to Interface: The minimum distance of the wild-type residue to the SARS-CoV-2 S-protein as resolved in PDB 6vw1<sup>9</sup>. mCSM-PPI2 $\Delta\Delta G$ : The predicted $\Delta\Delta G$ in kcal/mol for the missense mutation calculated by mCSM-PPI2<sup>12</sup> with PDB 6vw1 as the model structure. Prediction: Our assessment of the effect of the variant on ACE2 binding to SARS-CoV-2 S-protein on the basis of whether predicted $\Delta\Delta G$ exceeds our calibration thresholds (see Methods) and our structural assessment of the modelled variant. Mutagenesis: previously reported ACE2 mutant SARS-CoV S binding assay<sup>10</sup> that occur at the same position as the variant.*

#### 2.2.1 p.Gly326Glu is predicted to enhance ACE2-S interaction

p.Gly326Glu is predicted to increase the affinity of ACE2 for SARS-CoV-2 S by 1.0 kcal/mol. Figure 3 compares the ACE2-S interface in the vicinity of Gly326 with the mCSM-PPI2^12^ modelled p.Gly326Glu. Wild type Gly326 does not form any interactions with SARS-CoV-2 S, the closest approach between the proteins is 5.5 Å between ACE2 Gly326 Cα and the main-chain carbonyl of SARS-CoV-2 Thr500. The mutant p.Gly326Glu fills the void between the proteins and forms multiple H-bonding, polar and ionic interactions with the SARS-CoV-2 S backbone and residues Asn506 and Arg439. These new interactions are strongly suggestive that the mutant would lead to enhanced binding. However, the relevance of the interaction with Arg439 is uncertain because this residue was the result of a mutation (Asn439Arg) from the SARS-CoV-2 S reference (GenBank MN908947.1), engineered to enhance binding through interaction with ACE2 Glu329^9^. Modelling p.Gly326Glu in a structure with Asn at position 439 (6vw1-Asn439, Figure 3C, see Methods) yielded an identical predicted ΔΔG (i.e., 1.0 kcal/mol). In 6vw1-Asn439, the lost ionic interaction with the mutant Arg is compensated by an H-bond to the main chain amine of S Val503, which is accessible to a different rotamer conformation of the mutant Glu residue, whilst retaining the H-bond with S Asn506. As well as explaining why p.Gly326Glu is still predicted to enhance binding in the SARS-CoV-2 S model with Asn439, this highlights the sort of interplay that could occur between ACE2 variants and different SARS-CoV-2 strains. In this example of an engineered S-protein, the predicted effect on ΔΔG is the same for the ACE2 variant with both S-proteins but other SARS-CoV-2 S variants could interact differently with ACE2 variation.

**Figure 3.**
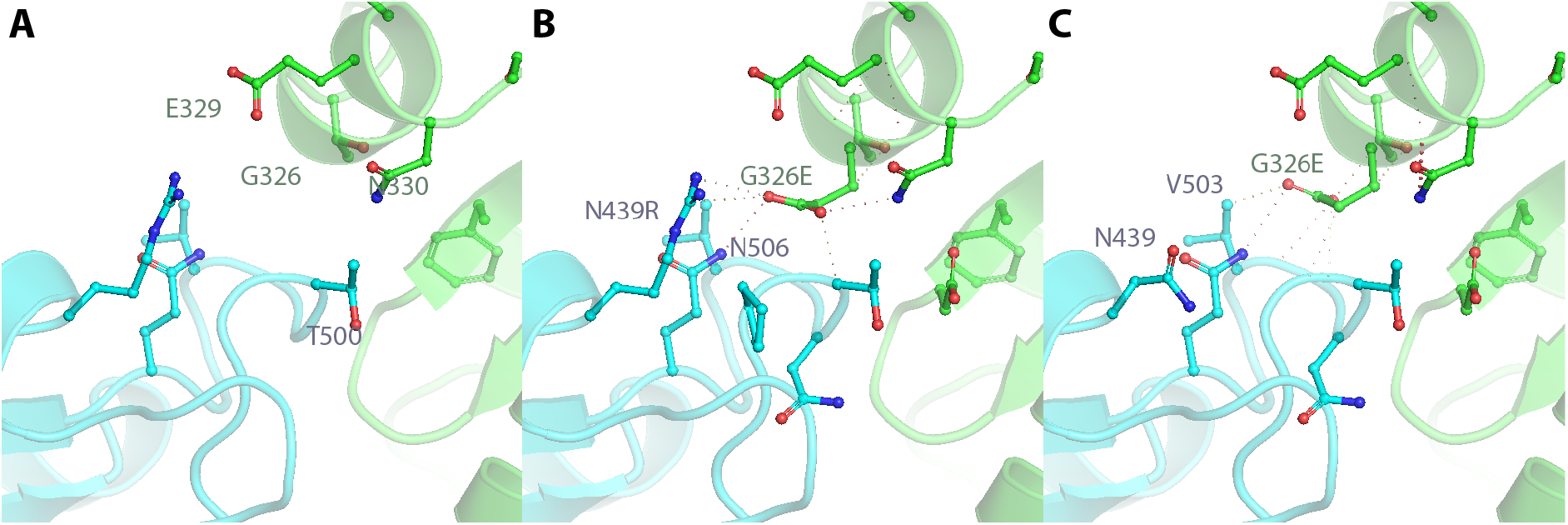
gnomAD^11^ ACE2 variant p.Gly326Glu gains new interactions with SARS-CoV-2 S that are likely to enhance binding. Local structural environments of A. ACE2 Gly326 in PDB 6vw1^9^ containing Asn439Arg in SARS-CoV-2 S; B. ACE2 p.Gly326Glu and C. ACE2 p.Gly326Glu in Asn439Arg reverse mutant model built with mCSM-PPI2^12^ from PDB ID: 6vw1. Dashed lines in B and C indicate H-bond, polar and/or ionic interactions. ACE2 backbone and side-chain carbons are coloured green and SARS-CoV-2 S is coloured blue.

#### 2.2.2 p.Asp355Asn, p.Glu37Lys and p.Gly352Val are predicted to inhibit ACE2-S binding

gnomAD ACE2 variants p.Asp355Asn, p.Glu37Lys and p.Gly352Val have mCSM-PPI2 ΔΔG < −1 kcal/mol and we therefore predict that they will significantly inhibit SARS-CoV-2 S binding or abolish it altogether. Figure 4 illustrates the structural differences between the wild-type and mutant structures of these mutants. p.Asp355Asn has the largest predicted negative ΔΔG of these mutations. Wild-type Asp355 is held in place by an intraprotein ionic interaction with Arg357. The loss of this charge complementarity in the p.Asp355Asn results in a different orientation for mutant Asn that introduces a number of clashes between this residue and others nearby, including the carbonyl oxygen of Thr500 of S. These clashes might be better accommodated in free ACE2 resulting in reduced affinity for the S-protein. In p.Glu37Lys, a favourable interprotein H-bond between wild-type Glu37 and RBD Tyr505 is lost alongside several intraprotein polar and ionic interactions. The new hydrophobic contact between the mutant Lys37 and the same Tyr residue is likely insufficient to compensate. p.Gly352Val is the farthest variant from the interface (5.4 Å) that is predicted to decrease S-protein binding and does not interact directly with any SARS-CoV-2 S residue. The effect on the interface is through a fourth-degree interaction that terminates with RBD Tyr505 (i.e., Gly352(C=O)-Tyr41-Lys353-RBD Tyr505). The introduction of Val into a position occupied by the buried Gly leads to several clashes. A few favourable internal hydrophobic contacts are also formed, including with Tyr41. The rearrangements necessary to accommodate Val might affect the bridged interaction between Tyr41 and RBD Tyr505 and the increased hydrophobic packing coupled with the loss of Gly could affect reduced flexibility. These considerations support the mCSM-PPI2 prediction.

**Figure 4.**
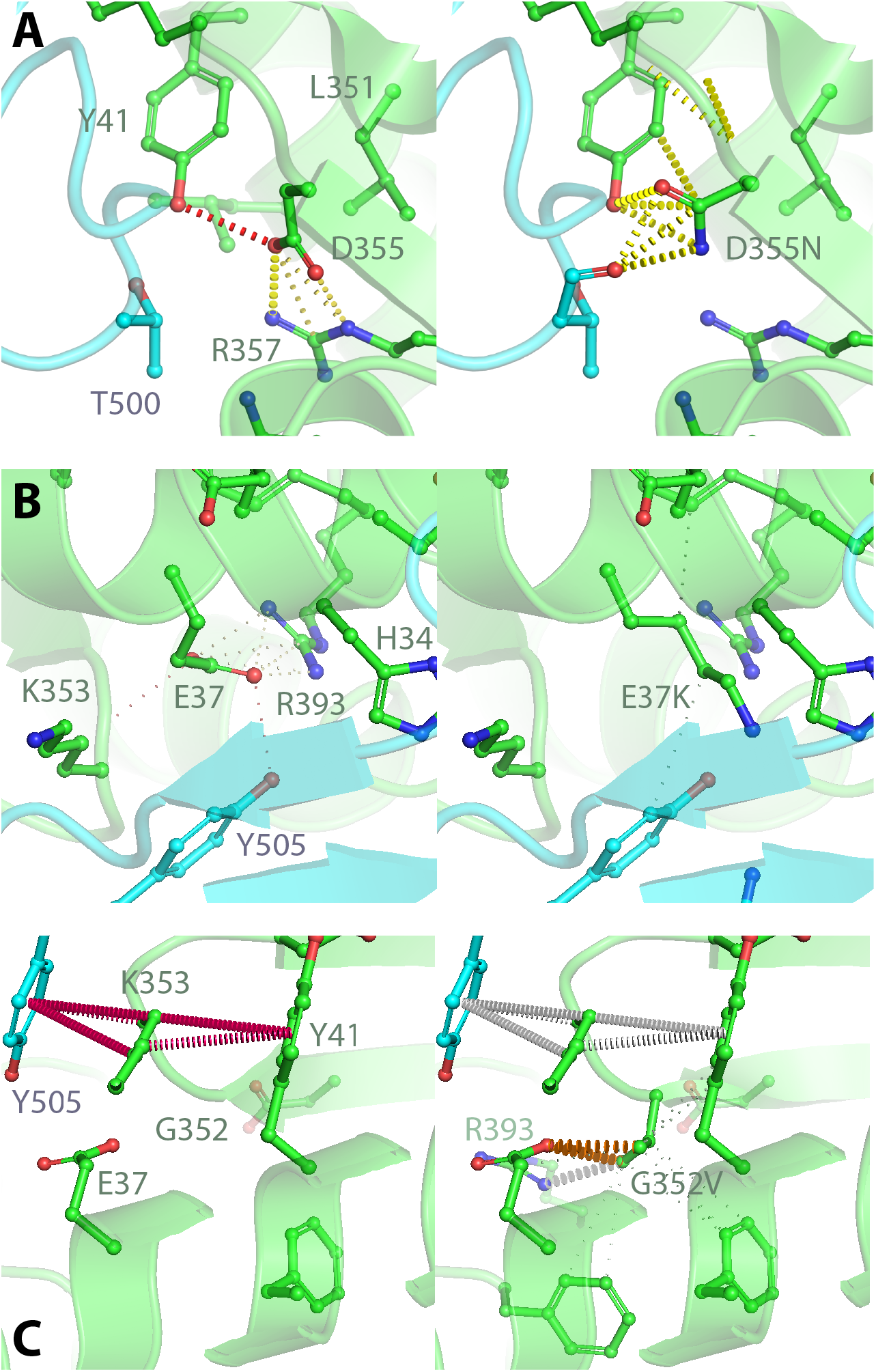
Structural comparison of ACE2 (green) gnomAD missense variants predicted to modify S-protein (light blue) interaction affinity. A. Increased clashes in the p.Asp355Asn mutant structure. Heavy dashed lines indicate atomic clashes. A H-bond/Polar clash is indicated by a red dashed line. B. Structural environment of p.Glu37Lys. C. Structural environment of p.Gly352Val. The wild-type structure is PDB ID: 6vw1^9^. The mutants were modelled onto 6vw1 with mCSM-PPI^12^. Figure created with PyMol^20^.

Mutation of residues on or near the interface are most likely to impact the interaction affinity but distant mutations can in principle affect interfaces through longer-range conformational effects. Figure 5 shows the distribution of predicted ΔΔG for 167 gnomAD ACE2 missense variants in residues resolved in PDB 6vw1. Whilst predicted ΔΔGs are biased toward a slight destabilising effect overall, the majority of variants have predicted ΔΔGs between −0.8 kcal/mol and 0.5 kcal/mol, suggesting that most have only a slight effect if any on ACE2-S binding. This is expected since most variants are away from the ACE2-S interface (Figure 5C) and distant conformational effects would be relatively rare anyway. In this case, no additional variants are found with ΔΔG < −1.0 kcal/mol, which is consistent with experimental evidence that ACE2-S binding is agnostic towards ACE2 conformational states^10^. These data also show that variants with predicted ΔΔG in excess of ±1 kcal/mol outlie the bulk of predictions, providing further empirical support for these thresholds.

**Figure 5.**
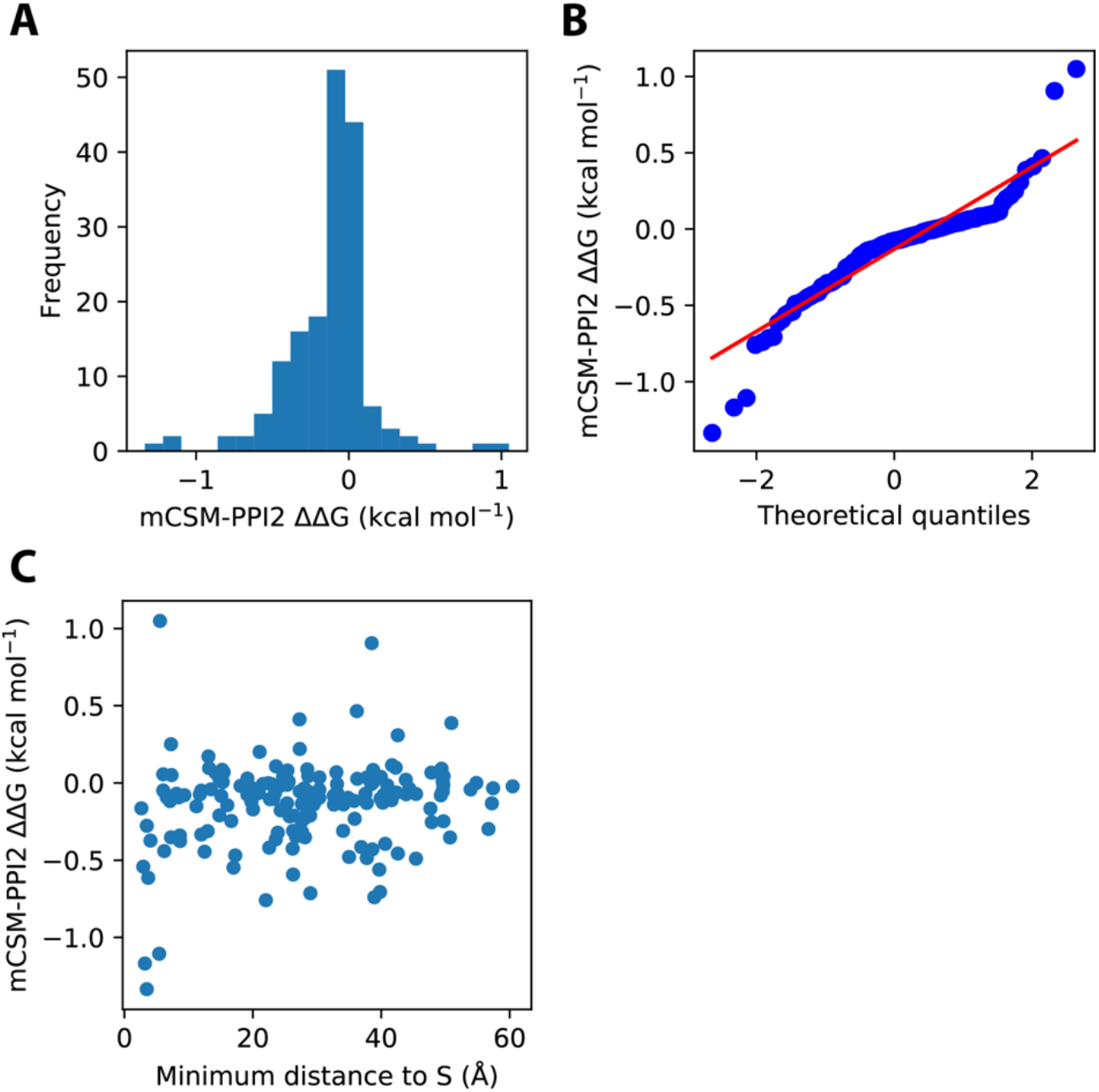
The distribution of mCSM-PPI2^12^ predicted ΔΔG for ACE2 gnomAD^11^ variants at residues resolved in PDB 6vw1^9^. A. Histogram of mCSM-PPI2 predicted ΔΔG. B. Probability plot of of mCSM-PPI2 predicted ΔΔG. C. mCSM-PPI2 predicted ΔΔG versus the distance of the variant site to the ACE2-S interface. Figure created with PyMol^20^.

There is a second outlier with relatively high positive ΔΔG that is highlighted by the probability plot (Figure 5B). This is p.Val447Phe with predicted ΔΔG = +0.9 kcal/mol, below the strict 1 kcal/mol threshold but well in excess of the relaxed threshold that we considered may be useful for potential ACE2-S affinity enhancing variants (see Methods). If this variant did lead to enhanced ACE2-S interaction, it could contribute significantly to the risk burden of ACE2 owing to its prevalence. In gnomAD this variant is observed in 13 individuals across three populations. This result is also unusual because the variant site is nearly 40Å from the ACE2-S interface. Figure 6 shows the local environment of Val447Phe.

**Figure 6.**
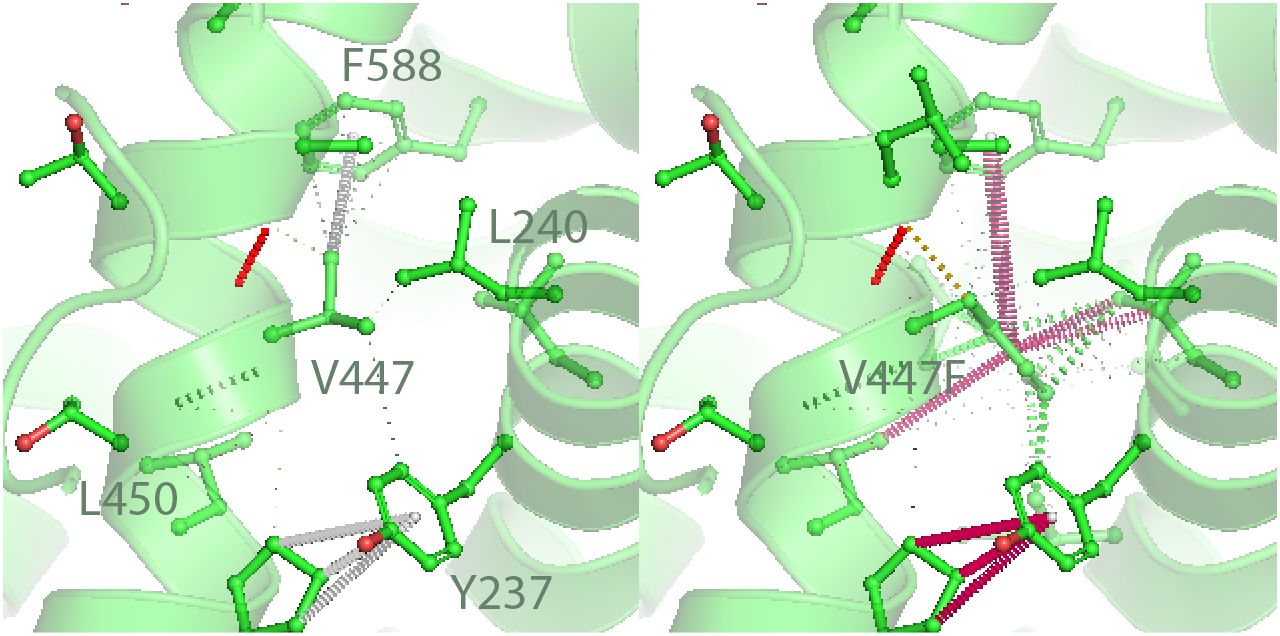
p.Val447phe: a second gnomAD^11^ variant with high predicted ΔΔG. Left: the local environment of ACE2 Val477 in PDB 6vw1. Right: mCSM-PPI2 provided model of p.Val447phe. Figure created with PyMol^20^.

The introduction of the larger Phe residue leads to a few clashes with neighbouring Ile233, Ile236 and Leu584. However, we were unable to discern a mechanism through which this mutant could modify ACE2-S binding as the mCSM-PPI2 mutant model showed no changes at the interface compared to wild-type. However, mCSM-PPI2 may recognise such motion in its internal model’s “atomic fluctuation” term^12^. However, in all the mutant models provided by mCSM-PPI2 we have evaluated, none display any relaxation of the structure, local or otherwise, in response to the mutant so we are unable to consider this factor in our critical evaluation. Due to the lack of structural corroboration of the mCSM-PPI2 predicted ΔΔG, we consider this variant to be of uncertain significance but worthy of follow up in cohort and experimental studies.

### 2.3 Exhaustive search for protective and risk factor ACE2 mutations

Existing human variation datasets are well-powered to detect common variation (1KG was estimated to detect >99% SNPs with MAF >1%^21^ and gnomAD is substantially larger) but they are far from comprehensive with respect to rare variation^11^. Rare variants with a significant effect on ACE2-S binding could have significant implications for the epidemiology of COVID-19 in addition to the consequences for affected individuals. If there were 10 such variants with an allele frequency of 1 in 50,000, their collective occurrence might be as high as 1 in 5,000 (discounting linkage) and when this is considered alongside the possibility that a high proportion of the global population will be exposed to SARS-CoV-2 it becomes clear that such effects should be investigated. In other words, the genetic burden of rare variants cannot be ignored.

#### 2.3.1 More mutations in ACE2 would reduce ACE2-S affinity than increase it

Figure 7 illustrates the distribution of ΔΔG predictions by mCSM-PPI2 from *in silico* saturation mutagenesis of the ACE2-S interface. Most of the 475 possible ACE2 mutations in residues at the interface with S are predicted to lead to a slight reduction in binding but there is a secondary mode < −1 kcal/mol containing 125 mutations that are predicted to lead to strongly reduced ACE2-S binding (Figure 7A). This constitutes a positive prediction rate of over 25% for inhibitory variants, a figure that is substantially higher than for the gnomAD variants (≈2%; §2.2) and the validation mutants (14%; Methods) due to the restriction of the analysis to sites at the ACE2-S interface where mutations are far more likely to effect the ACE2-S interaction. In contrast, only six mutations are predicted to result in enhanced binding at the strict ΔΔG threshold of > 1.0 kcal/mol for ACE2-S affinity enhancing variants and even at the relaxed threshold of > 0.5 kcal/mol only 14 mutations are highlighted. At face value, these results suggest that there is a good chance that individuals with ACE2 missense variants at these loci would have an ACE2 variant with reduced ACE2-S binding whilst ACE2 variants with enhanced affinity for S would be far less frequent.

**Figure 7.**
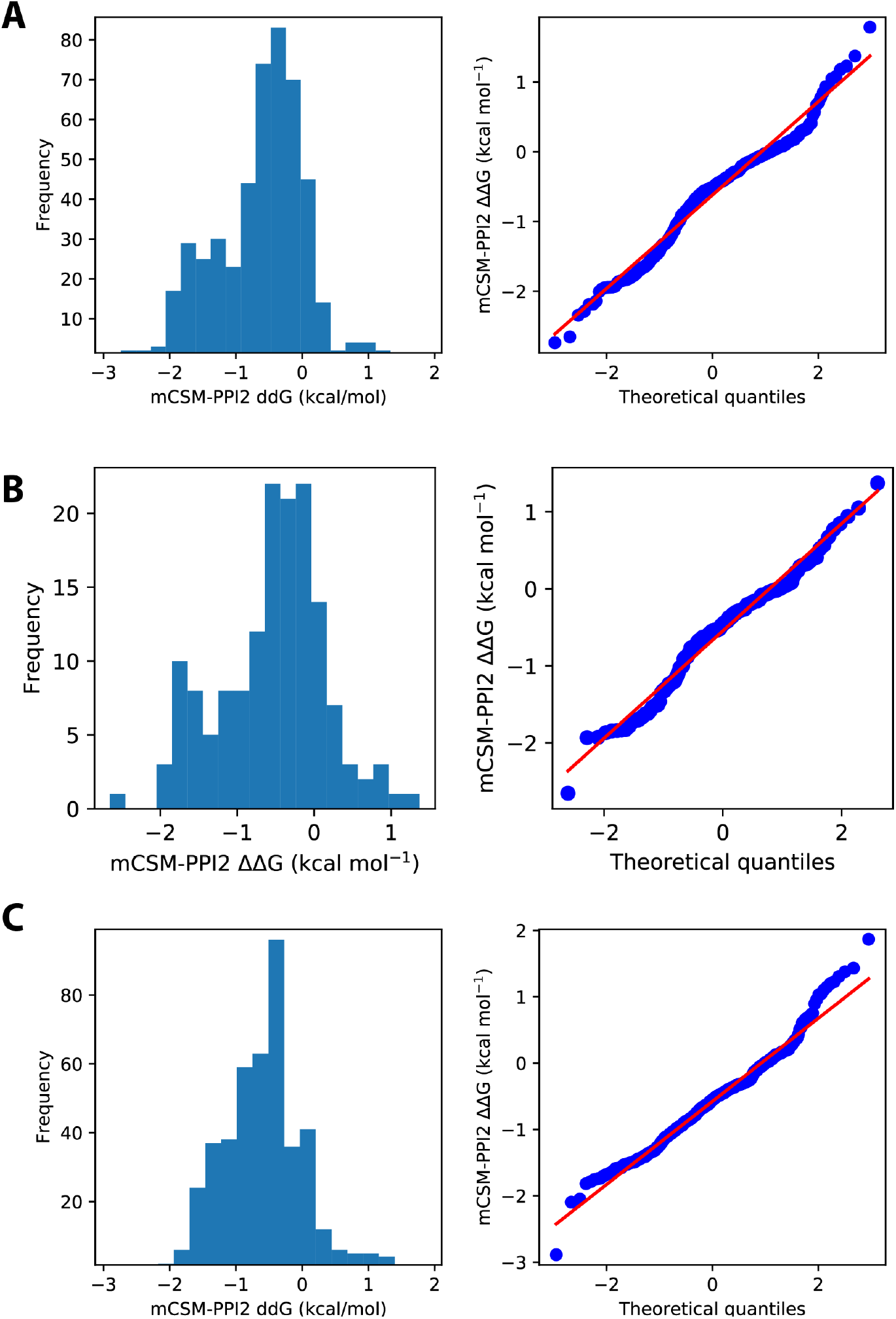
A. Distribution of mCSM-PPI2^12^ predicted ΔΔG from in silico saturation mutagenesis of the ACE2-S interface in PDB 6vw1^9^. A. predicted ΔΔG for 475 mutations across 25 sites on ACE2 corresponding to the 23 residues within 5 Å of SARS-CoV-2 S plus Gly326 and Gly352. B. predicted ΔΔG for the subset of 151 mutations across these sites that are accessible via a single base change of the ACE2 coding sequence. C. predicted ΔΔG for 437 mutations across 23 sites on SARS-CoV-2 S within 5 Å of ACE2.

Of the 475 possible mutations, the subset of 151 that are accessible via a single base change from the ACE2 coding sequence are more likely to be observed in the human population. Figure 7B shows the distribution of predicted ΔΔG for this subset. In this subset, there are 38 substitutions distributed over 12 sites with ΔΔG < −1.0 kcal/mol and a single substitution with ΔΔG > +1.0 kcal/mol is observed (Thr27Arg). Other factors can be considered too. For instance, any methylated CpG dinucleotides in the coding sequence would be particularly variable. This effect is observed at the protein level in the high mutability of Arg residues^22^. In ACE2, the only interface residue with a CG dinucleotide is Asp38. The residue is coded by GAC and the 3^rd^ base is part of a CG dinucleotide. Here the CG to TG mutation leads to a synonymous change but if we consider that this mutation might recur it is worthwhile assessing accessible missense changes from this codon, in this case, GAT to GGT (Gly) or GCT (Ala) are substitutions with ΔΔG < −1 kcal/mol. This approach could be applied to include other common variant codons such as those from high frequency synonymous variants. Another factor that might be relevant is the number of degenerate variants that lead to a given substitution.

This interpretation hinges on mCSM-PPI2 providing predicted ΔΔG that is unbiased. It is conceivable that mCSM-PPI2 could be biased towards negative ΔΔG predictions because the training datasets are skewed towards these observations^12,23^. Fortunately, the mCSM-PPI2 training data were augmented to mitigate this possibility^12^ and our own analyses of this bias in the ACE2 SARS-CoV S complex suggests that although the magnitudes of positive predictions can be depressed, overall the predictions are well-balanced (see Methods). In addition, some further evidence that mCSM-PPI2 ΔΔG is unbiased comes from the equivalent distribution of predicted ΔΔG for all possible mutations in SARS-CoV-2 S at sites that interact directly with ACE2 (Figure 7C). The ΔΔG distribution for these SARS-CoV-2 S mutations are still skewed towards ΔΔG < 0 kcal/mol but it does not contain the secondary mode below −1 kcal/mol that we observe in the equivalent ACE2 data (compare Figure 7A and C). Compared to ACE2, SARS-CoV-2 S also has a few more mutations that are predicted to enhance ACE2-S binding, including 10 > 1.0 kcal/mol out of 22 > 0.5 kcal/mol.

#### 2.3.2 A missense variant causing Thr27Arg would enhance ACE2-S affinity

We noted that one of the ACE2 mutations predicted to enhance ACE2-S binding was accessible from a single base change. This was Thr27Arg, which constitutes a second potential risk factor variant and deserves further evaluation. Thr27Arg is accessible from the C>G transversion in the wild-type codon ACA results in the AGA codon for Arg. Thr27Arg has the largest predicted ΔΔG of all the ACE2 binding-site mutants tested (1.4 kcal/mol) and is an apparent outlier amongst the full set of ACE2 saturation mutants (Figure 7A). Figure 8 illustrates the structural differences between the mutant and reference residue. Thr27 makes no specific contacts with SARS-CoV-2 S, coming closest to S Tyr489 (4.3 Å). Thr27Arg is amply accommodated in the interface. The guanidium interacts with S Tyr421 and Tyr473 whilst the charge is neutralised by nearby ACE2 Asp30. The guanidium fills a space between S Tyr473 and ACE2 Glu23 at the rim of the interface (behind the plane of Figure 8). The additional positive charge is neutralised by interaction with ACE2 Asp30. These features of Thr27Arg are strongly supportive of its potential to enhance ACE2-S binding.

**Figure 8.**
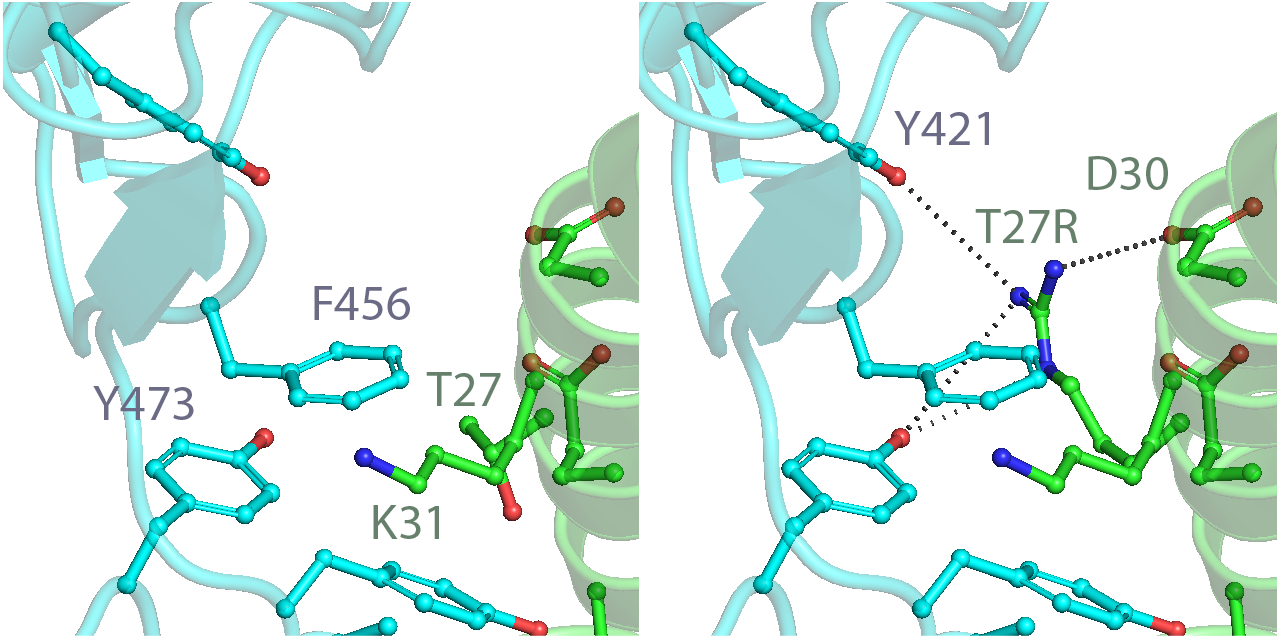
p.Thr27Arg: a second potential genetic risk factor missense variant in ACE2. The local environments of ACE2 Thr27 (left; PDB 6vw1^9^) and mutant p.Thr27Arg (right; mCSM-PPI2^12^ model derived from 6vw1). ACE2 (green; PDB chain A) and S-protein (light-blue; chain B). Dashed lines indicate side-chain interactions with the mutant Arg guanidinium group.

## 3 Other ACE2 variants and SARS-CoV-2 viral susceptibility

### 3.1 Missense variants in ADAM17 and TMPRSS2 cleavage regions

ACE2 can be cleaved by TMPRSS2 and ADAM17 and it has been shown that cleavage by TMPRSS2 enhances SARS-CoV cell entry^7^. Missense variants in the ADAM17 and TMPRSS2 cleavage regions (Table 4 and Figure 1) may affect the ratio of these ACE2 cleavage processes and consequently the availability of the TMPRSS2 ACE2 cleavage S-driven augmented entry pathway^7^. If this pathway were to prove active in SARS-CoV-2, then we could expect variants that disrupt ADAM17 cleavage to enhance SARS-CoV-2 infection whilst variants that disrupt TMPRSS2 might be protective. Mutations of Arg and Lys residues in the TMPRSS2 (R697-R716) and ADAM17 (R652-K659) cleavage regions can abolish ACE2 processing by these enzymes^7^. There are five Arg substitutions in the TMPRSS2 cleavage region that could reduce TMPRSS2 processing of ACE2 and consequently confer some level of resistance to SARS-CoV-2 infection, whilst the gain of Arg mutation p.Ser709Arg could lead to increased susceptibility by increasing TMPRSS2 processing. p.Arg697Gly is particularly notable given its prevalence and apparent specificity to South Asian males (Table 4). ADAM17 contains one Arg missense in gnomAD but since the variant leads to a Lys its effect in this respect is most likely neutral.

**Table 4.** Frequency of ACE2 gnomAD^11^ missense variants in the ADAM17 and TMPRSS2 cleavage regions.

| ID | Mutation | Allele Count | AC Male | AC Popn. Max | Popn. Max |
| --- | --- | --- | --- | --- | --- |
| <u>ADAM17 cleavage region</u> |  |  |  |  |  |
| - | p.Arg652Lys | 1 | 0 | 1 | NFE |
| - | p.Val658Ile | 1 | 0 | 1 | EAS |
| <u>TMPRSS2 cleavage region</u> |  |  |  |  |  |
| rs751603885 | p.Arg697Gly | 46 | 35 | 45 | SAS |
| rs780128908 | p.Ala703Ser | 3 | 0 | 3 | NFE |
| rs372986872 | p.Met706Ile | 1 | 0 | 1 | NFE |
| rs769062069 | p.Arg708Gln | 1 | 1 | 1 | NFE |
| rs776995986 | p.Arg708Trp | 1 | 0 | 1 | NFE |
| - | p.Ser709Arg | 1 | 0 | 1 | NFE |
| rs370187012 | p.Arg710His | 7 | 2 | 2 | EAS |
| - | p.Arg710Cys | 3 | 0 | 1 | EAS |

Column legend – Popn. Max: gnomAD population with highest allele frequency: non-Finnish European (NFE), East Asian (EAS) and South Asian (SAS).

### 3.2 Nonsense variants in ACE2

A few nonsense variants with potentially severe consequences for the canonical ACE2 transcript are present in gnomAD (Table 5). There are three stop-gained variants, two frameshift variants and a start-lost variant. Each of these variants would have a severe impact on the protein through the translation of a truncated and aberrant product or nonsense mediated decay. It is possible that stop-gained p.Leu656Ter and frameshifting p.Lys702ArgTer16 could lead to a semi functional product since they allow translation of a complete peptidase domain but the relevance to SARS-CoV-2 entry is unclear. An inframe deletion variant that results in the deletion of Lys313 occurs in a single male in the non-Finnish European cohort. It is difficult to interpret the effect of this variant as it could alter the pattern of exposed residues on the helix in which it occurs. However, we think it likely that it would be structurally accommodated, particularly since the closest loop begins only a few residues away at Gly319 (Figure 1). Crucially, Lys313 is about 20 Å from the SARS-CoV-2 RBD so we do not expect it to impact ACE2-S formation.

**Table 5.** Frequency of ACE2 gnomAD^11^ nonsense variants.

| ID | Mutation | Allele Count | AC Male | AC Popn. Max | Popn. Max |
| --- | --- | --- | --- | --- | --- |
| rs768110251 | p.Met1? | 1 | 0 | 1 | ASJ |
| - | p.Leu116Ter | 1 | 0 | 1 | ASJ |
| - | p.Lys313del | 1 | 1 | 1 | NFE |
| - | p.Gly422ValfsTer15 | 1 | 0 | 1 | NFE |
| - | p.Arg514Ter | 1 | 0 | 1 | FIN |
| rs199951323 | p.Leu656Ter | 1 | 0 | 1 | EAS |
| - | p.Lys702ArgfsTer16 | 0 | 0 | 0 |  |

Column legend – Popn. Max: gnomAD population with highest allele frequency:

## 4 Therapeutic applications

ACE2 has been identified as a potential therapeutic target for COVID-19^24^. One idea is to disrupt the ACE2-S interface with small-molecule or peptide inhibitors. The descriptions of the ACE2 variants provided here can help to anticipate possible pharmacogenomic effects of variants located at the ACE2-S interface. Another promising treatment direction is the administration of recombinant human ACE2 (rhACE2)^24,25^. This might compete with endogenous ACE2 for SARS-CoV-2 S. The approach has been proved-in-principle through the inhibition of SARS-CoV-2 infections in engineered human tissue models^25^ and a clinical trial is now underway (EudraCT Number: 2020-001172-15). An earlier trial was registered (Clinicaltrials.gov #NCT04287686)^24^ but has since been withdrawn before enrolling participants. The affinity of the administered rhACE2 might be a crucial component of the treatments efficacy, as indicated by the fact that mouse ACE2 did not have the same effect as human ACE2^25^. We will now describe how the ACE2-S interface saturation mutagenesis data can be used to design mutant ACE2 with tailored SARS-CoV-2 S affinity.

### 4.1 Design of therapeutic recombinant ACE2 with tailored affinity for SARS-CoV-2 S

In addition to predicting the effects of missense variants, the mCSM-PPI2 saturation mutagenesis data can be used to select substitutions in ACE2 that are likely to lead to reduced or enhanced affinity towards SARS-CoV-2 S. An ACE2 mutant with enhanced S affinity might be desirable for rhACE2 therapy. Even slightly enhanced affinity over endogenous ACE2 might shift the equilibrium towards rhACE2-S-protein complex formation and reduce the dose required for inhibition of viral infection. Table 6 presents the top mutations from the set of mutants predicted to lead to the largest increase in ΔΔG per site. These mutations are excellent candidates to improve the efficacy of rhACE2 therapy in COVID-19. We draw special attention to Gly326Glu, which corresponds to one of the gnomAD missense variants discussed earlier (see §2.2.1) and may be an excellent candidate since native ACE2 activity is likely to be maintained given that the variant is observed in a population dataset. The full dataset is provided in Supplementary Data File 1.

**Table 6.** ACE2 mutations predicted to significantly enhance affinity for SARS-CoV-2 S.

| Mutation | Distance to Interface | mCSM-PPI2 $\Delta\Delta G$ |
| --- | --- | --- |
| <u>$\Delta\Delta G > 1.0$ kcal/mol</u> |  |  |
| Thr27Arg | 3.7 | 1.4 |
| Phe28Trp | 3.4 | 1.2 |
| Leu79Trp | 3.5 | 1.2 |
| Gly326Glu | 5.5 | 1.0 |
| Gly352Tyr | 5.4 | 1.8 |
| <u>$\Delta\Delta G > 0.5$ kcal/mol</u> |  |  |
| Asp30Glu | 4.1 | 0.6 |
| Lys31Glu | 2.9 | 0.9 |
| Tyr41Asn | 2.6 | 0.8 |
| Gln42Glu | 3.1 | 0.8 |
| Leu45Glu | 3.9 | 0.9 |
| Tyr83Phe | 2.9 | 0.7 |
| Asn330Lys | 3.6 | 0.5 |
| Lys353His | 2.8 | 0.7 |

Column legend – Distance to Interface: The minimum distance of the wild-type residue to the SARS-CoV-2 S-protein as resolved in PDB 6vw1<sup>9</sup>. mCSM-PPI2 $\Delta\Delta G$ : The predicted $\Delta\Delta G$ in kcal/mol for the mutation calculated by mCSM-PPI2<sup>12</sup> with PDB 6vw1 as the model structure.

There are other criteria that should be considered to minimise the chance of disrupting the native activity of ACE2, which is thought could mitigate COVID-19 lung pathologies in addition to viral inhibition^24^. Basic considerations such as avoiding mutations to or from proline or glycine and nuanced strategies such as restricting candidates to sites with known population variants are relevant. The former criterion might be ignored if a major physicochemical change is deemed necessary to enhance S affinity. Whether or not this interface has any endogenous role will underlie tolerance of native ACE2 activity to mutations.

The utility of an ACE2 variant with enhanced ACE2-S binding as a competitive inhibitor of endogenous ACE2 is clear, but since our data could be used to guide the design of a variant with depressed binding, we can consider if this might be of any use. Assuming that rhACE2 therapy works by competing with membrane bound native ACE2 for SARS-CoV-2 S (rhACE2 efficacy in acute respiratory distress syndrome is thought to be due to circulating ACE2 activity^26^, i.e. it does not enter the cell), an ACE2 mutant with reduced affinity for SARS-CoV-2 could be beneficial if it replaced native ACE2 on the membrane. This could be useful if gene therapy were practical or desirable for this disease.

As far as we know, the clinical trial assessing the efficacy of rhACE2 therapy in COVID-19 uses recombinantly expressed wild-type cleaved ACE2^25^. Clinical testing and approval can be expediated for repurposed existing therapies as is clearly the strategy here. Our proposal to for enhanced rhACE2 could in principle be achieved with a single mutation and might be suitable for delivery within a similar rapid development framework.

## 5 Discussion and conclusion

### 5.1 The impact and prevalence of ACE2 variant receptor activity genotypes

How far could ACE2-S affinity modulation variants effect SARS-CoV-2 infection? There is clear evidence that mutations in ACE2 that effect its affinity for coronavirus S-protein could provide complete protection from infection. This can be understood from the host range specificity of coronaviruses driven by ACE2-S complementarity alongside mutagenesis studies showing that controlled single point mutations can abolish interaction^10^. In SARS-CoV-2, it has been shown that the APN and DDP4 cell entry pathways are not active^1,5^, so there can be no recourse to these routes (unless mutations in these genes could establish this activity). In this work, some observed ACE2 variants were predicted to have a comparable effect on ACE2-S binding as mutants that were experimentally determined to abolish the interaction altogether. Therefore, it is possible that individuals carry ACE2 variants that confer resistance to SARS-CoV-2 infection. If infection efficiency operates on a continuous scale, perhaps to some extent proportional to the binding affinity, then various gradations of resistance could be conferred by different variants.

It is less clear what effect the ACE2-S binding enhancer variants might have. In principle, they could increase susceptibility to infection by promoting SARS-CoV-2 cellular attachment, potentially reducing the infective dose of SARS-CoV-2. ACE2 variants with enhanced receptor activity could also promote infection of cell types that express lower levels of ACE2, and so lead to increased risk of COVID-19 complications (e.g., acute renal failure) since viral spreading is associated with clinical deterioration^27^. Such effects might also lead to increased transmission from individuals with ACE2-S enhancing variants. It was suggested that enhanced ACE2-S affinity in SARS-CoV-2 versus SARS-CoV could lead to increased infection in the upper respiratory tract^5^, this hypothesis is beginning to bear out with evidence of upper respiratory tract infection^28^ and higher ACE2 affinity of SARS-CoV-2 compared to SARS-CoV now established^9^. However, there is also the possibility that such variants are protective, if for instance the ACE2 variant led to a persistent stable ACE2-S complex that interfered with S proteolysis, although to our knowledge this phenomenon has not been observed. Experimental work-up of the Thr27Arg and Gly326Glu variants of ACE2 should provide answers to these questions.

Aside from ACE2-S affinity, host cell ACE2 expression levels are known to be important for the cellular specificity of SARS-CoV^29^ and attention has been directed toward ACE2 expression levels as a potential factor in COVID-19 susceptibility and severity^30^. It is helpful to consider that with respect to SARS-CoV-2 cell entry, the physical mechanism underlying the effects of variation in ACE2 expression and ACE2 variant S-protein affinity is the formation of the ACE2-S complex. In a simplistic model, ACE2 expression modifies ACE2 membrane concentration whilst ACE2-S affinity modifies the equilibrium dissociation constant. It is conceivable then that affinity variants and expression variants compound the effect of one another, in a manner akin to that suggested for ACE2 variants and ACE2 stimulating drugs^30^, and so ACE2 haplotypes and linkage with other regulatory loci could be critical. This also has implications for more moderate ACE2-S affinity modifying variants, since their effect could be amplified in the presence of altered ACE2 expression.

Other epidemiological features can be reconciled with our observations. For instance, males are significantly more likely to die as a result of COVID-19 infection. Since ACE2 resides on the X chromosome, female heterozygotes will express a proportion of ACE2 alleles whilst males are hemizygotes and carry only a single ACE2 allele (this was also noted by Asselta et al.^31^ who also noted that ACE2 escapes complete X-inactivation). A male carrying a rare ACE2-S affinity enhancing variant will be worse affected than a female heterozygote. Moreover, it is conceivable that in females the more resistant ACE2 allele could rapidly become predominant in infected tissues due to selection induced by host cell turnover over infection cycles, gradually increasing the expression of the S-protein resistant ACE2 allele if present. In addition to the benefits of this selection process, this may have implications for female survivors of COVID-19 depending on the persistence X-inactivation bias and the nature of the hitchhiking alleles.

Although, as described above, there is clear evidence that missense variants are capable of abolishing ACE2’s receptor activity towards SARS-CoV-2 S, it is complicated to estimate the prevalence of such variants in the population. The gnomAD variants predicted to reduce ACE2-S binding are all very rare. p.Glu37Lys is the most prevalent with an allele frequency (AF) 0.0003 in the gnomAD Finnish samples and it is also observed in one African/African American sample. The two other gnomAD inhibitory variants p.Gly352Val and p.Asp355Asn are both doubletons observed only in non-Finnish Europeans, each corresponding to AF = 0.00003 in this population. Expanding on this, a simple estimation is that these two variants might confer SARS-CoV-2 resistance to just over 1 in 15,000 non-Finnish Europeans, comparable to many rare diseases. However, these are not the only potentially protective ACE2 variants. Our analysis of the effects of all possible missense mutations at the ACE2-S interface predicted that 38 substitutions accessible via single base changes at 12 sites would lead to severely reduced interaction. These substitutions account for 41 missense variants out of the possible 108 single nucleotide variants at the codons corresponding to the ACE2 residues that bind SARS-CoV-2 S. A crude upper bound under the highly conservative assumption that these variants occur at lower frequencies than the allele frequency of a singleton can be calculated. The minimum possible AF for a singleton in gnomAD non-Finnish Europeans is 8.8 × 10^−6^. Since we are calculating the effect of 41 unobserved variants, their joint frequency (assuming no linkage) is 3.6 × 10^−4^, which is equivalent to 72 variants per 200,000 alleles or 7.2 per 10,000 population. A more sophisticated estimate takes account of the empirical detection of rare variants in ACE2. For example, there are 1,181 variant alleles in 56,885 non-Finnish Europeans arising from variants with AF < 0.01 in the 2,415 nucleotide ACE2 coding sequence (n.b. at this AF threshold, 90 % are missense). This amounts to a variation rate of about 4.3 × 10^−6^ per nucleotide per allele. So, at the 12 sites (36 nucleotides) with variants predicted to strongly reduce ACE2-S binding we could roughly expect 1.6 × 10^−4^ variants per allele or 31 variants per 100,000 population. This is another conservative estimate for these sites since surface residues tend to be more variable than the core or other functional regions of the protein^32^. Finally, since 41 out of 108 possible SNPs at these 12 sites are predicted to lead to a strongly reduced ACE2-S interaction, we can multiply by this proportion to estimate the number that might strongly reduce ACE2-S binding. This yields an estimate that 11.8 individuals per 100,000 population possess an ACE2 allele that is predicted to be resistant to SARS-CoV-2 infection.

Although these rough estimates suggest that highly resistant ACE2 alleles are extremely rare it is important to note that allele frequencies show significant variation even between large variation datasets^33^. Moreover, it is known that local population allele frequencies also vary substantially from those reported in public datasets^34^ and it is therefore possible that some populations have resistance mutations at higher frequencies. Also, we have only considered ACE2 variants that are predicted to result in a strong reduction or total abolition of ACE2-S binding. Our data also suggests that ACE2 interface mutants tend to at least slightly reduce ACE2-S binding and this means that there will be a higher prevalence of ACE2 alleles with this property. In concert with altered ACE2 expression levels, these moderate ACE2-S affinity variants could have a larger effect at the population scale.

### 5.2 Applications to population genetic analyses of ACE2 variation in COVID-19

Another approach to assess the role of ACE2 variation in COVID-19 has been the application of comparative population genetic analysis. Two reports have found polymorphisms in ACE2 expression quantitative trait loci (eQTL) vary between populations but both concluded there was no clear evidence regarding ACE2 missense variants^4,31^. Why did neither of these studies find evidence that ACE2 missense variants could be associated with variable population specific epidemiology? First, Rosanna et al.^31^ acknowledge that there is no conclusive evidence that apparent cross-population differences in COVID-19 epidemiology are not accounted for by population demography^31^, which raises the possibility that population differences in allele frequencies and burden are meaningless in the context of COVID-19 genetics. Nevertheless, the detection of decreased TMPRSS2 burden in Italians and Europeans compared to East Asians^31^ showed that the approach could turn up interesting results with rational mechanistic interpretations. A potential prejudice against the resolution of COVID-19 genetic association with ACE2, and one that highlights an application of our work, is the inclusion criteria for variants in burden tests. Standard filtering strategies based on predicted variant severity may have worked well for TMPRSS2 where SARS-CoV-2 exploits its native catalytic function^5^, but ACE2 mediated SARS-CoV-2 cell entry does not require ACE2 catalytic activity^10^ and as a result these filters might be counterproductive. Inclusion criteria for missense variants might be better based on their predicted effect on S-protein affinity, differentiating variants predicted to reduce ACE2-S affinity from those that enhance it since they may have opposite effects (Table 3), or potential to mitigate some other SARS-CoV-2 infection pathway such as TMPRSS2 or ADAM17 cleavage region variants (Table 4). Moreover, it might be useful to test missense variants with specific effects on SARS-CoV-2 infection separately, to isolate these features from the other roles ACE2 may have in COVID-19 pathology. This application to burden tests would likely prove useful to attempts to determine host genetic factors that influence COVID-19 susceptibility and severity.

### 5.3 Reconciling conflicting results from closely related studies

Two studies applied computational methods to gnomAD variants at the ACE2-S interface and report conflicting conclusions to our own^35,36^. Hussain et al.^35^ employed homology modelling, pathogenicity predictions and the PRODIGY^37^ webserver to assess 17 gnomAD ACE2 variants located at the S interface or at reported mutagenesis sites, all of which were considered in our work, including variants p.Asp355Asn, p.Glu37Lys and p.Gly352Val that we find are likely to strongly inhibit ACE2-S binding (Table 3). They tentatively suggested that p.S19P and p.E329G might be resistant to S-protein attachment, which partially conflicts with our conclusion that these variants are unlikely to significantly effect ACE2-S binding (Table 3), and they did not predict a significant reduction in ACE2-S binding in the p.Asp355Asn, p.Glu37Lys and p.Gly352Val variants. In a different study, Othman et al.^36^ assessed eight gnomAD ACE2 variants at the interface. Their variant models were derived from homology models built from SARS-CoV S bound to human-civet chimeric ACE2 (PDB ID: 3scl) and they also used PRODIGY^37^ in addition to another predictor to calculate ΔΔG. After finding no variants exceeded a predefined 1 kcal/mol threshold they concluded that this meant there was little chance of ACE2 mediated resistance to SARS-CoV-2. Amongst the eight variants tested in Othman et al.^36^ were the three variants we predict will strongly disrupt ACE2-S binding and p.Gly326Glu, which we find enhances ACE2-S binding.

Although these negative findings conflict with our work, we remain confident in our results for a few reasons. First, we point to our validation data on mCSM-PPI2, which demonstrates its effectiveness in the highly similar SARS-CoV S-protein ACE2 complex (see Methods), and we report that PRODIGY, which was employed by both other studies, did not perform as well on this validation set (Supplementary Figure 2). Another factor that could contribute to the different results is that the other studies derived variant models from different structures of ACE2-S than we have (i.e., PDBs 6lzg^38^, and 3scl). We found mCSM-PPI2 predictions from PDBs 6vw1^9^ and 6lzg^38^ to be highly concordant for the same interface mutations (ρ = 0.97; see also Supplementary Figure 3) confirming that structure selection was not an issue.

With respect to p.Ser19Pro and p.Glu329Gly, which Hussain et al.^35^ predicted might be resistant to S-protein attachment on the basis of PRODIGY predicted ΔG and the presence of fewer charge interactions in the p.Glu329Gly model and other lost contacts. An interesting rationale given for these results was that the interaction between ACE2 Lys353 and the S-protein, a site shown to be important for binding SARS-CoV S-protein^10^, was lost in their models of p.Ser19Pro and p.Glu329Gly but it is unclear how confident this movement is given that Lys353 is about 30 Å and 16 Å away from the variant sites, respectively. In our results, p.Ser19Pro had the smallest predicted ΔΔG (−0.2 kcal/mol) of any of the gnomAD interface variants whilst p.Glu329Gly had the third smallest (−0.4 kcal/mol; Table 3). These values are well below our calibrated 1.0 kcal/mol threshold, but we do recognise the possibility they could be false negatives or could weakly inhibit ACE2-S binding. However, we observed nothing in these sites’ local environments that suggests the variant residues would strongly affect ACE2-S association (§2 and Supplementary Table 1) and since Hussain et al.^35^ refer to distant structural changes to explain the result we think that the mCSM-PPI2 result for these variants is more structurally realistic.

The difference between our conclusions and these reports highlights the uncertainty inherent in prediction methods and the variability in predictions from different algorithms and models, but we are confident that the results presented in our work are strongly justified. The mCSM-PPI2 prediction algorithm was designed specifically for the task in hand, to predict changes in protein binding affinities upon mutation, and displays best-in-class performance according to community best-practice benchmarks^12^. Our validation of mCSM-PPI2 with experimental data from the closely related SARS-CoV S-ACE2 complex provides direct evidence that this algorithm yields accurate predictions for this complex and calibrates the algorithm’s quantitative predictions with observed physical behaviour (Methods). Additionally, we have followed up key results with a critical structural inspection of the local environments of the mutant models and in our judgement the predictions are in line with the principles established by the original structural and mutagenesis work in this area^8,10^.

### 5.4 Conclusion

In summary, we have assessed the potential of ACE2 genetic variation to impact SARS-CoV-2 infection, primarily through prediction of ACE2-S binding efficiency, but also with respect to the potential ACE2 TMPRSS2 cleavage driven augmented entry pathway. Our results are based largely on predictions from the state-of-the-art mCSM-PPI2^12^ mutation effect predictor for protein-protein complex affinity, but we first validated this method against published experimental ACE2 mutant SARS-CoV S-protein affinities^10^ and demonstrated its applicability to this system and calibrated the quantitative predictions to physical observations (see Methods). We found a single known variant in the gnomAD^11^, p.Gly326Glu, that we predict enhances ACE2 binding affinity for SARS-CoV-2 S that is therefore a potential risk factor for COVID-19. We found three variants in gnomAD that are predicted to reduce ACE2 affinity towards S, p.Glu37Lys, p.Gly352Val and p.Asp355Asn, that are therefore potentially protective against COVID-19. The models that underlie these predictions were structurally evaluated and we judged the predictions to be in line with the established principles concerning ACE2 receptor recognition by coronavirus S-proteins^8–10,39^. We extended our predictions to all possible ACE2 missense variants that interact with SARS-CoV-2 S. We found that mutations that lead to reduced ACE2-S interaction affinity are far more likely than mutations that lead to increased interaction. Simple estimates of the prevalence of ACE2 resistance conveying alleles suggest that the most protective alleles will be very rare (12-70 per 100,000 population) and potential risk factors will be rarer but the possibility of higher prevalence in local or underrepresented populations remains. Also, weaker effects, which are more common, could be amplified by ACE2 expression polymorphisms. This is consistent with the high rate of asymptomatic and mild infections and the rarity of severe disease in low-risk groups. A further candidate risk factor variant was identified in this analysis, Thr27Arg. Finally, we described how the mCSM-PPI2 predicted mutant affinities could be used to enhance the potency of a recombinant ACE2 therapy that is currently undergoing clinical trial^24,25^ in COVID-19. This work has clear applications in helping to prioritise experimental work into SARS-CoV-2 S-protein human ACE2 recognition; developing genetic diagnostic risk profiling for COVID-19 susceptibility and severity; improving detection and interpretation in future COVID-19 genetic association studies and in developing therapeutic agents for COVID-19.

## Methods

### Integration of structure, variant and mutagenesis data

We conducted an initial scoping analysis of ACE2 in Jalview 2.11^14^. The ACE2 protein sequence was retrieved from UniProt^13^ and the structure of SARS-CoV-2 S-protein bound ACE2 (PDB ID: 6vw1) was mapped to this sequence with Jalview’s built-in structure mapping capability. The structure was visualised in a Jalview^14^ linked UCSF Chimera^16^ session. Contacts between ACE2 and SARS-CoV-2 S-protein were identified with the Chimera FindClash command with default settings for contacts, and Van der Waals overlaps were stored as attributes. The average overlap per residue was pulled into Jalview as features from Chimera with the Fetch Chimera attributes function. This allowed UniProt features (e.g., mutagenesis, SARS-CoV binding, TMPRSS2 cleavage site) to be interactively inspected alongside the residue S-protein contacts and gnomAD variants loaded from a Jalview feature file (Supplementary Figure 1; see below).

The pyDRSASP suite^15^ was used to integrate 3D structure, population variant and mutagenesis assay data for analysis. Population variants from gnomAD were mapped to ACE2 with VarAlign^15^. Residue mappings were derived from the Ensembl VEP annotations present in the gnomAD VCF. In addition, we manually checked the gnomAD multi nucleotide polymorphisms (MNPs) data file and found no records for ACE2.

ACE2 mutagenesis annotations were retrieved from UniProt with ProteoFAV^15^. Li *et al.*^10^ is indicated as the source for all SARS-CoV Spike-ACE2 mutagenesis annotations. The UniProt/Swiss-Prot annotations included a qualitative description of the effect of each mutant. These descriptions were parsed into terms and represented with ordinal categories: Abolishes interaction > Strongly inhibits > Inhibits > Slightly inhibits > No effect. Quantitative results were extracted from figures in Li *et al.*^10^ with WebPlotDigitizer 4.2^40^. We validated the data transcription by comparing the tabulated values with the UniProt qualitative annotations (Supplementary Figure 4). We transcribed data for additional mutants from Li *et al.*^10^ that were not reported in UniProt.

Structural data were retrieved with ProIntVar^15^. The structure of chimeric SARS-CoV-2 Spike receptor binding domain in complex with human ACE2 (PDB ID: 6vw1)^9^ was downloaded from PDBe. Residue-residue contacts were calculated with ARPEGGIO^18^. ACE2-S interface residues were defined as those with any interprotein interatomic contact (Supplementary Table 1). The asymmetric unit contains two ACE2-S-protein dimers. 68 residue contact pairs were observed in both complexes in the asymmetric unit, 6 were unique to complex A-E and 4 were unique to complex B-F. Although the complex between chain A ACE2 and chain E SARS-CoV-2 S is assigned as the most probable biological unit by RCSB and PDBe, for the purposes of completeness, we counted all contacts in both complexes as the ACE2-S interface. Interface contact numbers are defined as the number of residues on the partner protein a residue is in contact with (i.e., we do not count intraprotein contacts). For example, ACE2 S19 contacts SARS-CoV-2 S residues Ala475, Gly476 and Ser477 so its interface contact number is three. Solvent accessibility was calculated with DSSP^41^ for the intact complex and each individual PDB chain (corresponding to monomeric ACE2 and SARS-CoV-2 S) and relative solvent accessibilities (RSA) were derived using the maximal allowed residue accessibility^42^. Interface residues were classed as Core or Rim as suggested by David and Sternberg^17^ on the basis of differences between RSA of the complex and free monomers. UniProt-PDB mappings were retrieved from SIFTS^43^.

### Prediction of missense variant effects on S-protein – ACE2 interaction

The mCSM-PPI2^12^ web server was used to predict the effect of mutations on the SARS-CoV-2 Spike-ACE2 interface topology and binding affinity with the structure PDB ID: 6vw1^9^. ACE2 gnomAD variants were supplied to the web server via upload of a mutation list to the Predictions interface. Wild-type and mutant interaction graphs were inspected online with the Single-Point Mutation interface. The wild-type and mutant models were downloaded from the server as PyMol sessions and detailed structural comparisons were conducted with PyMol^20^. Our typical workflow was to group all the objects in each session (i.e., wild-type and mutant) and then load both models and contact descriptor objects into a single PyMol session. The models within the mCSM-PPI2 PyMol sessions did not retain their original numbering and it was necessary to reload the original PDB 6vw1 model to identify residues conveniently.

Most of the saturation mutagenesis data were obtained by selecting the Saturation Mutagenesis option on the Interface Analysis submission page. In this analysis, the mCSM-PPI2 web server defines the interaction interface automatically. No residue farther than 5 Å from the interface site, as defined by the web server, was included in the analysis. Saturation mutagenesis data for positions Gly326 and Glu352, which were beyond this limit but found in our study to harbour ACE2-S affinity altering variants, were obtained via the mutation list submission. Data were downloaded in CSV format. n.b. The mCSM-PPI2 model for p.Gly352Tyr displayed a chemically invalid conformation for the mutant Tyr and so results for this mutant were excluded.

mCSM-PPI2 predictions for mutations corresponding to the ACE2 single-point mutants with experimental data^10^ on SARS-CoV Spike-ACE2 binding were obtained with the structure of SARS-CoV S receptor binding domain complexed with human ACE2 (PDB ID: 2ajf)^8^ as starting model. These results were merged with the experimental binding assay data, collected as outlined above.

mCSM-PPI2 predictions were obtained for models with the engineered Arg439 substituted with the native Asn. Arg439Asn was modelled with mCSM-PPI2 on the structure 6vw1. The model was exported to PDB from the PyMol session downloaded from the mCSM-PPI2 results page. This model was provided as the starting structure for mCSM-PPI2 predictions. mCSM-PPI2 predictions for the validation reverse mutations were obtained in the same way.

### Validating mCSM-PPI2 Spike-ACE2 ΔΔG predictions on SARS-CoV S-protein data

mCSM-PPI2^12^ is a random forest family predictor that models mutations with graph based signatures and other descriptors trained on protein-protein binding affinities from the SKEMPI 2.0 database^23^. mCSM-PPI2 predicts the change in the Gibbs free energy of the complex as a result of point mutations (ΔΔG). Negative ΔΔG means that the mutation is predicted to decrease the binding affinity whilst positive ΔΔG predicts an increase in affinity. mCSM-PPI2 achieved an average Pearson’s correlation (ρ) of 0.82 and root mean square error (RMSE) of 1.18 kcal/mol during 10-fold cross-validation and ranked first out of 26 others on the CAPRI^44^ community standard benchmark. A better representative of the predictor’s generality to different complexes is the cross-validation results on a training set with low redundancy of complexes. This gave ρ = 0.75 and RMSE = 1.30 kcal/mol. This behaviour is typical for machine learning methods where performance is best in systems closer to the training data. The method was also evaluated on a set of 378 alanine scanning mutations across 19 protein complexes and this resulted in exceptional cross-validation metrics ρ = 0.95 and RMSE = 0.25 kcal/mol, which can be interpreted as an indication of the consistency of the method for different mutations within complexes

Experimentally determined SARS-CoV S-protein affinities are available for several ACE2 mutants^10^. These mutants present the opportunity to validate mCSM-PPI2 predictions on a system closely related to SARS-CoV-2 S-protein interactions. If mCSM-PPI2 can be shown to perform well here, then we expect good performance for SARS-CoV-2 S-protein affinities, potentially approaching the accuracy displayed in the evaluation of the alanine scan dataset. Methods Figure 1A illustrates the performance of mCSM-PPI2 ΔΔG as a predictor for the experimentally determined S-protein interactions. The experimental affinities are positively correlated with predicted ΔΔG (ρ = 0.68) so that mutants in ACE2 that strongly reduced or abolished SARS-CoV S interaction (S−) tend to have lower predicted ΔΔG than mutants with little or no effect on the interaction (S+). In detail, five of the seven S-mutations have predicted ΔΔG between −2 and −1 kcal/mol whilst most S+ mutations have predicted ΔΔG > −0.5 kcal/mol. The remaining two S-mutants, Lys31Asp and Lys353His, received ΔΔG predictions > −0.5 kcal/mol, which is within the range displayed by the bulk of S+ mutants. Two S+ mutants received slightly lower ΔΔG predictions than most other S+ mutants, but still well above −1 kcal/mol. These results suggest that ΔΔG < −1.0 kcal/mol yields a confident prediction that the mutation strongly inhibits or abolishes ACE2-S interaction (rule-in threshold) whilst ΔΔG > −0.5 kcal/mol is indicative that the mutation has little to no effect on the interaction (rule-out threshold). ΔΔG predictions between these limits are relatively sparse but S+ mutations received predicted ΔΔG as low as −0.73 kcal/mol whilst the first inhibitory mutant had predicted ΔΔG −1.17 kcal/mol. Table 7 shows the concordance between the experimental result and the mCSM-PPI2 prediction setting a prediction threshold at ΔΔG < −1.0 kcal/mol. At this threshold, no S+ mutants are falsely predicted to abolish ACE2-S binding and only two S-mutants are falsely predicted to have little to no effect on the interaction.

**Methods Figure 1.**
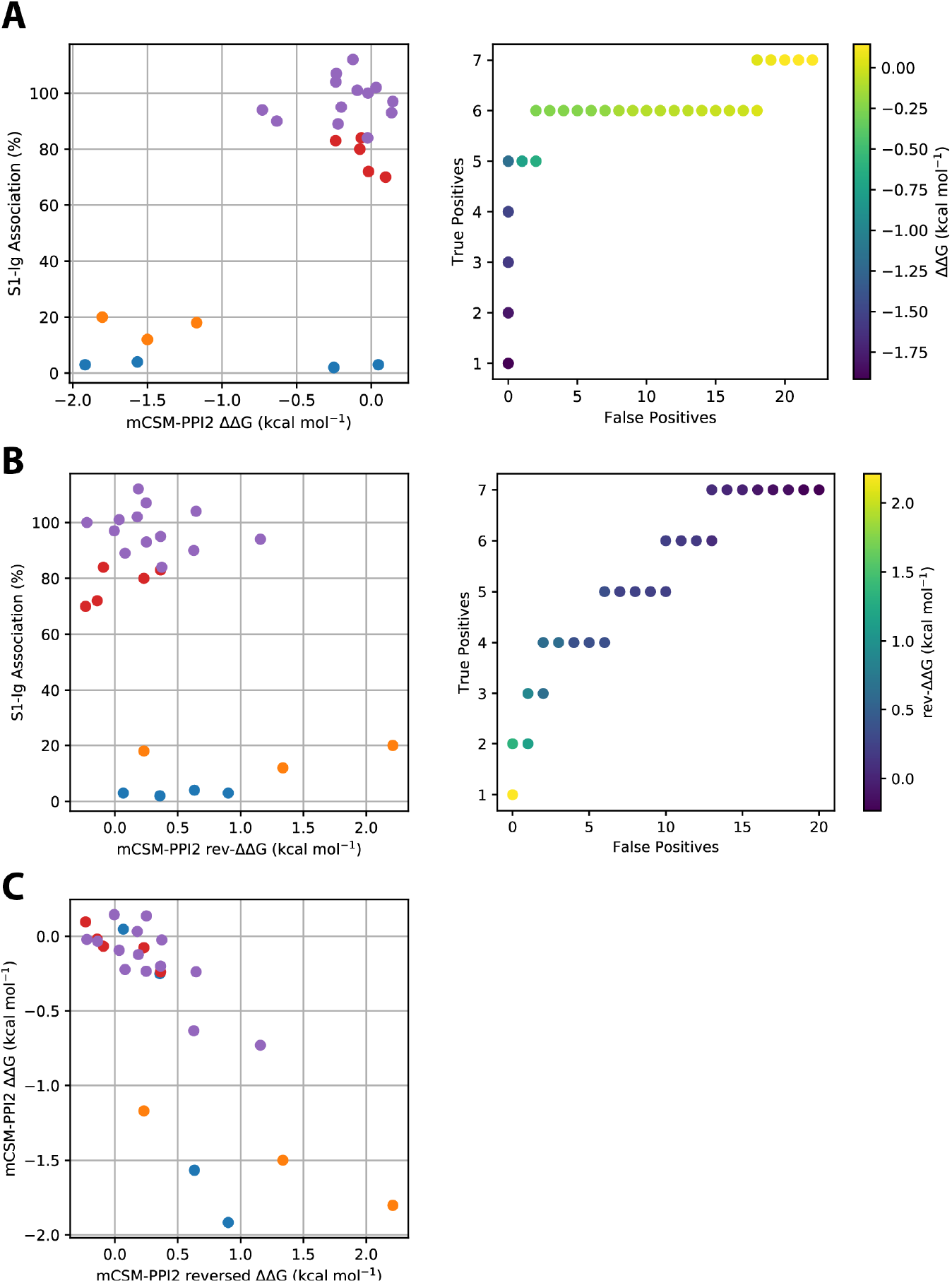
Validation of mCSM-PPI2^12^ predicted ΔΔG with reported ACE2 mutants with SARS-CoV S binding data^10^ modelled in PDB 2ajf^8^. A. Left: Relative proportion of human ACE2 immunoprecipitated with the S1 domain of SARS-CoV S^10^ versus mCSM-PPI2 predicted ΔΔG. Mutants are coloured according to the UniProt description of the effect on SARS-CoV binding: no effect (red), slightly reduced binding (green), strongly reduced binding (orange) and abolished interaction (blue). Right: ROC-like curve illustrating performance of mCSM-PPI2 ΔΔG as a predictor of ACE2-S binding. Mutants that strongly inhibited or abolished the ACE2-S interaction are considered positive cases (S+) whilst mutants that only slightly inhibited or had no effect on the interaction are considered negative cases (S−). B. Validation of mCSM-PPI2^12^ capacity to identify ACE2-S affinity enhancing variants. Left: Relative proportion of ACE2 immunoprecipitated with the S1 domain of SARS-CoV S^10^ versus mCSM-PPI2 predicted ΔΔG^reverse^ (e.g., ΔΔG^reverse^ for Asp355Asn is the predicted ΔΔG for Asn355Asp in the Asp355Asn model structure; see Methods). Mutants are coloured as in A. Right: ROC-like curve illustrating performance of mCSM-PPI2 ΔΔG^reverse^ as a predictor of ACE2-S binding. Definition of true positives and false positives as in A. C. Comparison of mCSM-PPI2 ΔΔG with ΔΔG^reverse^, i.e., predicted ΔΔG for the reverse mutation. See Supplementary Table 3 for data values.

**Table 7.**
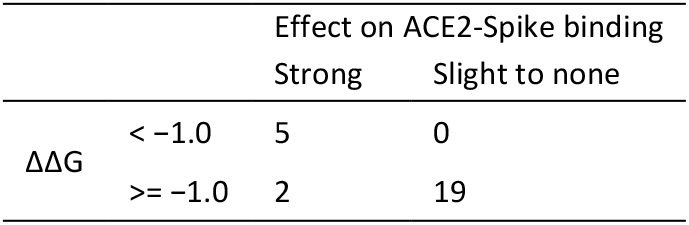
Prediction of ACE2 mutant SARS-CoV S binding classification based on mCSM-PPI2^12^ predicted ΔΔG modelled with PDB 2ajf^8^. Mutants are predicted to lead to a strong to total reduction in ACE2-S interaction if mCSM-PPI2 ΔΔG < −1.0 kcal/mol and have little to no effect otherwise. Experimentally positive cases are mutants described as abolishing or effecting a strong reduction in ACE2-S interaction (S− mutants) whilst negative cases are those described as having little to no effect (S+ mutants).

We can evaluate the mutant models to try to explain why two of the validation mutants are misclassified. The discordance between the mCSM-PPI2 prediction and experiment for Lys353His is surprising because mCSM-PPI2 correctly identifies the other Lys353 mutants, Lys353Ala and Lys353Asp, as causing a severe reduction in the ACE2-S interaction. Methods Figure 2 compares the interactions of the wild-type and mutant complexes at these sites. The size of the residues is a clear difference between the mutants, resulting in fewer hydrophobic contacts in the Lys353Ala and Lys353Asp mutants. Additionally, Lys353His can form ring-ring interactions with Tyr759 on the RBD, which could compensate for the fewer hydrophobic contacts in this mutant. These considerations go some way to explain why Lys353His is predicted to have less impact on ACE2-S binding than Lys353Ala or Lys353Asp, but they do not explain the discrepancy with the experimental result. One factor that might be at play is that the protonation state of the mutant His could modulate its inhibition of ACE2 S binding and this might not be well-modelled by mCSM-PPI2. The second misclassified mutation is another Lys mutant. Methods Figure 3 shows the environment of ACE2 Lys31 and Lys31Asp. The sidechain of Lys31 is oriented away from the SARS-CoV S interface allowing the Lys β- and ɣ-carbons to form hydrophobic interactions with S Tyr442 and Tyr475. In free ACE2, Lys31 is oriented outward and is fully exposed and it has been suggested that the internal neutralisation of Glu35 by Lys31 in the ACE2-S complex is required for efficient binding^39^. In Lys31Asp, there are fewer hydrophobic interactions, but the carboxyl group forms an H-bond with Tyr442 that could compensate. These interactions might explain the relatively small predicted ΔΔG of −0.25 kcal/mol and it could be that mCSM-PPI2 does not recognise the importance of the internal neutralisation for binding.

**Methods Figure 2.**
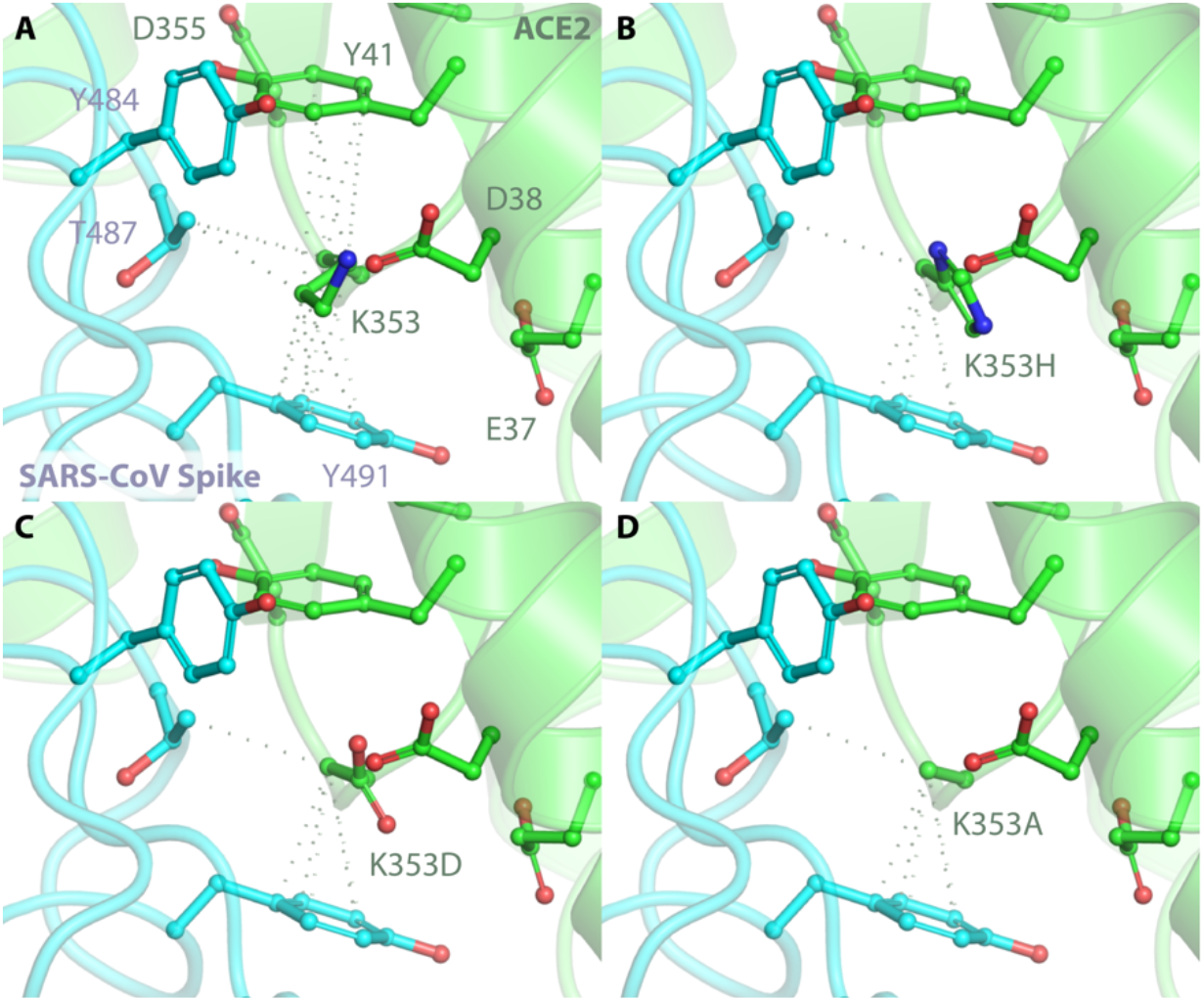
ACE2 Lys353 mutants receive different mCSM-PPI2 ΔΔG predictions for different residue substitutions. The local environments of A. human ACE2 Lys353 (PDB ID: 2ajf^8^), B. Lys353His, C. Lys353Asp and D. Lys353Ala (B, C and D: mCSM-PPI2 model from PDB 2ajf). Human ACE2 is coloured green whilst SARS-CoV S is blue. Dashed lines indicate hydrophobic interactions as labelled by mCSM-PPI2. Figure created with PyMol^20^.

**Methods Figure 3.**
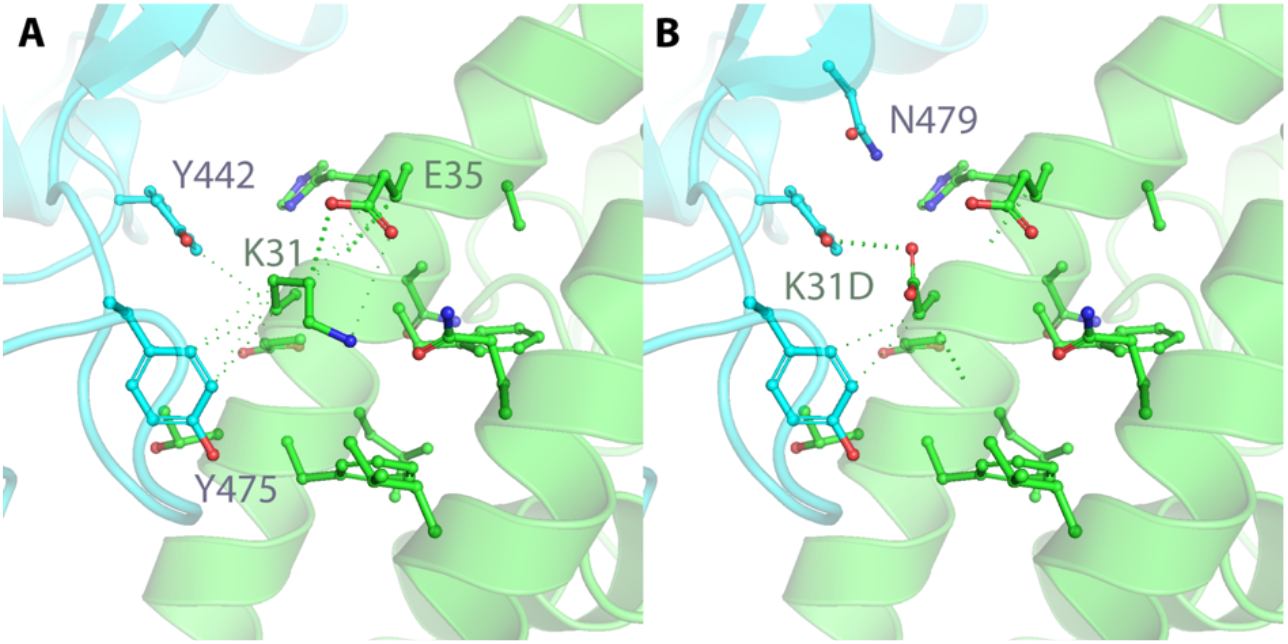
Lys31Asp is misclassified as having little effect on ACE2-S interaction by mCSM-PPI2^12^. The local environments of A. human ACE2 Lys31 (PDB ID: 2ajf^8^) and B. Lys31Asp (mCSM-PPI2 model from PDB 2ajf^8^). Human ACE2 is coloured green whilst SARS-CoV S is blue. Dashed lines indicate hydrophobic and H-bond interactions. Figure created with PyMol^20^.

Our validation set does not include any mutants predicted to enhance ACE2-S binding. The potential candidates we found were mutant ACE2 orthologs with humanising mutations^10^ but we were unable to find a suitable experimentally determined protein structure to model these mutations. In the absence of these data we are unable to validate or calibrate mCSM-PPI2 explicitly for ACE2 variants that are predicted to enhance S binding but there are a few options to address this deficit. First, we recognise that the ability to predict mutations that increase ΔG as well as those that lower ΔG was part of the design specification for mCSM-PPI2^12^. This objective was advanced by extending the mutant affinity training data with hypothetical reverse mutations to mitigate the bias towards destabilising mutations^12^. With this in mind, we could simply accept this stated capability of the algorithm at face value and assume a symmetric threshold of +1.0 kcal/mol to identify mutations that will significantly enhance binding. This is technically equivalent to extrapolating the relationship between ΔΔG and experimental ACE2-S association (Methods Figure 1A) past 0 kcal/mol, which warrants some explanation. Forward extrapolation of the linear trend in Methods Figure 1A above 0 kcal/mol would yield predicted assay results > 100 %, reaching ≈ 200 % by +2.0 kcal/mol. Although values greater than 100% were observed in the experimental binding assay (and are valid)^10^, the assay is bounded above by an unknown quantity that in our estimation should correspond to the inverse of the ratio of immunoprecipitated ACE2 by the C-terminal tag antibody to the total amount of ACE2 in the sample. This ceiling represents the point at which all mutant ACE2 is immunoprecipitated by the S1-Ig fusion protein used in the original assay^10^.

An alternative approach is hinted at by the reverse mutations the mCSM-PPI2 developers used to augment their training dataset with complex stabilising mutations. Since ΔΔG is a thermodynamic state function, ΔΔG^reverse^ = −ΔΔG^forward^, and any deviation in the predicted result is an error^12^. This means that the reverse mutations of the experimental mutants that inhibit binding lead to enhanced binding. These can be readily modelled, for example, to model the reverse mutation for Asp355Asn we submitted the mCSM-PPI2 model of Asp355Asn and requested a prediction for Asn355Asp. Methods Figure 1B illustrates the relationship between the experimental binding assays and predicted ΔΔG^reverse^. Generally, ΔΔG^reverse^ follows our expectations (Methods Figure 1C), with most ΔΔG^reverse^ > 0 kcal/mol and displaying a negative linear correlation with ΔG^forward^ (ρ = −0.78), but there are a few moderate outliers amongst the S- mutations with apparently underestimated ΔΔG^reverse^. These outliers contribute to a reduction in correlation with the experimental assays compared to that achieved with the forward mutations (Methods Figure 1B, ρ = 0.41), which leads to poorer performance as a predictor of the assay results (Methods Figure 1B). An overall reduction in concordance was expected, since starting from predicted models runs the risk of compounding errors, but it is not clear if the reduced sensitivity is due to generally poorer sensitivity of mCSM-PPI2 towards mutations that enhance affinity or a consequence of these errors. However, we can still achieve good specificity with ΔΔG thresholds of > 1.0 kcal/mol (TPR = 0.29, TNR = 0.95) or slightly more relaxed > 0.5 kcal/mol (TPR = 0.57, TNR = 0.84).

These results indicate that mCSM-PPI2 is applicable to the ACE2 coronavirus S complex and that ΔΔG is a good predictor for S-protein affinity. A threshold of ΔΔG < −1.0 kcal/mol is calibrated to a strong reduction or total abolition of ACE2 S binding affinity. For potential affinity enhancing variants, ΔΔG > +1.0 kcal/mol is a sensible starting point that should yield high specificity but if increased sensitivity is desirable then a relaxed threshold as low as ΔΔG > + 0.5 kcal/mol is appropriate.

### mCSM-PPI2 predictions are robust to the presence of engineered Arg439 in PDB 6vw1

The structure of the ACE2 SARS-CoV-2 S-protein receptor binding domain (RBD) complex contains a chimeric SARS-CoV-2 S-protein with the engineered substitution S Asn439Arg introduced to increase binding with ACE2. mCSM-PPI2 predicts that S Asn439Arg enhances S-protein since mutations to the wild-type is predicted to destabilise ACE2-S (ΔΔG = −0.25 kcal/mol for S Arg439Asn in PDB 6vw1; ΔΔG^reverse^ = 0.11 kcal/mol), although notably this prediction does not meet our strict thresholds. Although the mutation itself does enhance ACE2-S binding^10^, its presence does not appear to significantly influence mCSM-PPI2 predicted ΔΔG of gnomAD variants. This is shown in Methods Figure 4, which compares predictions from the chimeric RBD (i.e., PDB 6vw1) to those from a model of the RBD with Asn439 modelled in 6vw1; these are practically identical (ρ = 1.00). Despite this, the variant residue conformations can be influenced by the presence of the engineered residue (§2.2.1) but it appears that for all gnomAD variants, where this happens the alternative conformations yield the same or very similar ΔΔG.

**Methods Figure 4.**
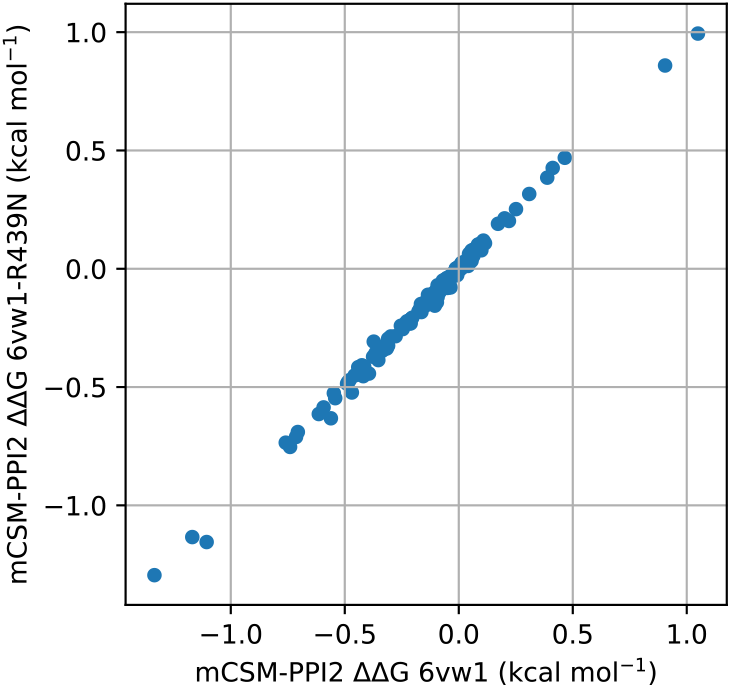
Comparison of mCSM-PPI2^12^ predicted ΔΔG for all ACE2 gnomAD^11^ variants that map to PDB 6vw1^8^ calculated from PDB 6vw1 directly and from the mCSM-PPI2 model of R439N in PDB 6vw1.

### Enumerating possible ACE2 missense SNPs

The ACE2 gene (ENSG00000130234) was retrieved from Ensembl in Jalview^14^. Two identical CDS transcripts (ENST00000427411 and ENST00000252519) were found with the Get Cross-References command corresponding to ACE2 full-length proteins (ENSP00000389326 and ENSP00000252519). These correspond to the UniProt ACE2 sequence Q9BYF1. The CDS was saved in Fasta format and this was parsed in Python with Biopython. The CDS was broken into codons and all possible single base changes were enumerated and translated using the standard genetic code. This provided the set of amino acids accessible to each residue via a single base change.

### Software

Jalview 2.11^14^ was used for interactive sequence data retrieval, sequence analysis, structure data analysis and figure generation. UCSF Chimera^16^ and PyMol^20^ were used for structure analysis and figure generation.

The pyDRSASP^15^ packages, comprising ProteoFAV, ProIntVar and VarAlign were used extensively for data retrieval and analysis as described in Methods. Biopython was used to process sequence data.

Data analyses were coded in Python in Jupyter Notebooks. Numpy, Pandas and Scipy were used for data analysis. Matplotlib and Seaborn were used to plot data.

## Code availability

All code and analysis notebooks used in this study are available from the Barton Group public GitHub repository at https://github.com/bartongroup/covid19-ace2-variants.

## Data availability

This study employed several public datasets, which are available from their original sources and all necessary identifiers and accessions are provided in the methods. Data that was compiled and calculated as part of this study, including all mutant models of ACE2 gnomAD variants and ACE2 validation mutants, are available from the Dundee Resource for Sequence Analysis and Structure Prediction COVID-19 portal accessible from http://www.compbio.dundee.ac.uk/drsasp.html.

## Acknowledgements

We thank Dr Jim Procter for alerting us to the release of the structure of ACE2 in complex with SARS-CoV-2 S and his initial observations regarding variants at the ACE2-S interface. We thank the Dundee Research Computing team for supporting our IT infrastructure and remote working. This work was supported by Biotechnology and Biological Sciences Research Council Grants (BB/J019364/1 and BB/R014752/1) and Wellcome Trust Biomedical Resources Grant (101651/Z/13/Z).

## Author contributions

S.A.M performed the research, compiled and analysed data, and drafted the manuscript. G.J.B and S.A.M conceived the research, interpreted results and edited and reviewed the final manuscript.

## Author information

G.J.B and S.A.M: Division of Computational Biology, College of Life Sciences, University of Dundee, UK

## Extended Data

**Supplementary Table 1.** Structural description of the ACE2 – Spike-protein interface.

| Residue | Contact Number | Min. VdW Distance | S-protein counterparts | ARPEGGIO Interaction Types |
| --- | --- | --- | --- | --- |
| S19 | 3 | 0.0 | A475, G476, S477 | Polar-Bond, VdW-Bond, Clash |
| Q24 | 6 | 0.0 | A475, N487, G476, F486, S477, Y489 | H-Bond, Clash |
| T27 | 5 | 0.4 | A475, N487, F456, Y473, Y489 | VdW-Proximal |
| F28 | 1 | 0.2 | Y489 | H-Bond, Polar-Bond |
| D30 | 2 | 0.7 | L455, F456 | Hydrophobic-Bond |
| K31 | 6 | 0.0 | Q493, L455, L492, F456, F490, Y489 | Hydrophobic-Bond, Clash |
| H34 | 3 | 0.0 | Q493, L455, Y453 | Aromatic-Bond, H-Bond, Hydrophobic-Bond, Polar-Bond, VdW-Bond, Clash |
| E35 | 1 | 0.0 | Q493 | H-Bond, Polar-Bond, Clash |
| E37 | 1 | 0.1 | Y505 | H-Bond, Hydrophobic-Bond, Polar-Bond |
| D38 | 4 | 0.0 | Q498, G496, HOH701, Y449 | H-Bond, Polar-Bond, VdW-Bond, Clash |
| Y41 | 4 | 0.0 | N501, Q498, HOH701, T500 | H-Bond, Polar-Bond, VdW-Bond, Clash |
| Q42 | 4 | 0.0 | Q498, HOH701, T446, Y449 | VdW-Bond, Clash |
| L45 | 2 | 0.4 | Q498, T500 | VdW-Proximal |
| L79 | 2 | 0.1 | G485, F486 | Hydrophobic-Bond |
| M82 | 1 | 0.1 | F486 | Hydrophobic-Bond, VdW-Bond |
| Y83 | 3 | 0.0 | N487, F486, Y489 | Aromatic-Bond, H-Bond, Hydrophobic-Bond, Polar-Bond, VdW-Bond, Clash |
| Q325 | 2 | 1.4 | R439, V503 | VdW-Proximal |
| E329 | 1 | 1.0 | R439 | VdW-Proximal |
| N330 | 1 | 0.3 | T500 | VdW-Proximal |
| K353 | 7 | 0.0 | N501, Q498, G496, G502, HOH701, F497, Y505 | H-Bond, Hydrophobic-Bond, Polar-Bond, VdW-Bond, Clash |
| G354 | 3 | 0.2 | N501, G502, Y505 | Polar-Bond |
| D355 | 2 | 0.2 | G502, T500 | VdW-Proximal |
| R357 | 1 | 0.2 | T500 | VdW-Proximal |
| R393 | 1 | 0.5 | Y505 | VdW-Proximal |
| N90-glycan (BMA703) | 2 | 0.1 | R408, T415 | H-Bond, Polar-Bond, VdW-Bond |

**Supplementary Table 2.** mCSM-PPI2 predictions for all gnomAD ACE2 missense variants in residues resolved in PDB 6vw1.

| ID | Mutation | Distance to Interface | mCSM-PPI2 $\Delta\Delta G$ | Allele Count | AC Male | AC Popn. Max | Popn. Max |
| --- | --- | --- | --- | --- | --- | --- | --- |
| rs73635825 | p.Ser19Pro | 2.6 | -0.2 | 40 | 6 | 39 | AFR |
| - | p.Ile21Thr | 7.3 | 0.3 | 1 | 1 | 1 | AMR |
| rs778030746 | p.Ile21Val | 7.3 | -0.4 | 2 | 2 | 2 | NFE |
| rs756231991 | p.Glu23Lys | 6.2 | -0.4 | 1 | 0 | 1 | NFE |
| rs4646116 | p.Lys26Arg | 6.0 | 0.0 | 704 | 258 | 85 | ASJ |
| - | p.Lys26Glu | 6.0 | 0.1 | 1 | 1 | 1 | NFE |
| rs781255386 | p.Thr27Ala | 3.7 | -0.6 | 2 | 0 | 2 | AMR |
| - | p.Glu35Lys | 2.9 | -0.5 | 3 | 1 | 2 | EAS |
| rs146676783 | p.Glu37Lys | 3.2 | -1.2 | 6 | 3 | 5 | FIN |
| - | p.Phe40Leu | 8.6 | -0.3 | 3 | 0 | 3 | AMR |
| - | p.Ser43Arg | 8.5 | -0.1 | 1 | 0 | 1 | SAS |
| - | p.Tyr50Phe | 13.0 | -0.3 | 1 | 0 | 1 | NFE |
| rs760159085 | p.Asn51Asp | 14.2 | 0.1 | 2 | 1 | 2 | SAS |
| rs775273812 | p.Thr55Ala | 15.3 | 0.0 | 1 | 1 | 1 | NFE |
| rs771621249 | p.Asn58Lys | 12.0 | -0.3 | 2 | 0 | 2 | SAS |
| - | p.Asn58His | 12.0 | -0.1 | 2 | 1 | 2 | NFE |
| rs759162332 | p.Gln60Arg | 15.5 | 0.1 | 2 | 0 | 2 | NFE |
| - | p.Met62Val | 13.5 | 0.0 | 1 | 1 | 1 | NFE |
| - | p.Asn64Lys | 13.2 | 0.1 | 1 | 1 | 1 | AFR |
| rs755691167 | p.Lys68Glu | 7.3 | 0.1 | 2 | 2 | 2 | SAS |
| - | p.Phe72Val | 8.7 | -0.4 | 1 | 1 | 1 | EAS |
| - | p.Ser77Phe | 9.3 | -0.1 | 1 | 1 | 1 | NFE |
| rs766996587 | p.Met82Ile | 3.5 | -0.3 | 0 | 0 | 0 |  |
| rs766996587 | p.Met82Ile | 3.5 | -0.3 | 2 | 0 | 2 | AFR |
| rs759134032 | p.Pro84Thr | 7.1 | -0.1 | 1 | 0 | 1 | SAS |
| rs746808776 | p.Gln86Arg | 13.1 | 0.2 | 2 | 1 | 2 | NFE |
| rs763395248 | p.Thr92Ile | 15.1 | -0.1 | 2 | 2 | 2 | NFE |
| - | p.Gln102Pro | 14.8 | -0.2 | 2 | 0 | 1 | NFE |
| rs143158922 | p.Asn103His | 14.8 | 0.0 | 2 | 0 | 2 | AFR |
| rs139773121 | p.Val107Ala | 19.8 | -0.1 | 2 | 1 | 2 | AFR |
| rs201900069 | p.Arg115Gln | 30.4 | 0.0 | 30 | 15 | 9 | EAS |
| rs751227277 | p.Ser128Thr | 28.6 | -0.1 | 1 | 0 | 1 | SAS |
| rs766124365 | p.Pro138Ala | 45.4 | -0.1 | 1 | 1 | 1 | NFE |
| - | p.Cys141Tyr | 38.5 | 0.0 | 1 | 0 | 1 | AFR |
| - | p.Asn154Lys | 38.4 | 0.0 | 2 | 0 | 1 | AMR |
| rs746034076 | p.Asn159Ser | 49.4 | 0.1 | 2 | 1 | 1 | AMR |
| - | p.Ala164Ser | 42.6 | 0.3 | 1 | 0 | 1 | AFR |
| rs769593006 | p.Glu166Gln | 45.4 | -0.5 | 1 | 0 | 1 | NFE |
| rs748076875 | p.Glu171Val | 41.6 | -0.1 | 1 | 1 | 1 | NFE |
| rs754511501 | p.Gly173Ser | 40.6 | -0.4 | 4 | 2 | 1 | SAS |
| rs779651019 | p.Pro178Leu | 40.2 | -0.1 | 1 | 0 | 1 | NFE |
| rs758142853 | p.Val184Ala | 33.1 | -0.1 | 8 | 1 | 8 | EAS |
| rs750052167 | p.Leu186Ser | 28.7 | -0.1 | 1 | 0 | 1 | AMR |
| - | p.Met190Thr | 23.6 | 0.1 | 1 | 0 | 1 | SAS |
| rs765733397 | p.Ala191Pro | 24.7 | 0.0 | 1 | 0 | 1 | NFE |
| rs762219565 | p.Ala193Glu | 21.1 | 0.2 | 3 | 0 | 3 | AMR |
| rs764661406 | p.His195Asn | 19.1 | -0.1 | 1 | 1 | 1 | SAS |
| rs764661406 | p.His195Tyr | 19.1 | 0.0 | 1 | 0 | 1 | NFE |
| - | p.Glu197Gly | 28.0 | 0.0 | 2 | 0 | 1 | ASJ |
| - | p.Asp198Asn | 29.0 | -0.7 | 1 | 0 | 1 | NFE |
| rs750145841 | p.Tyr199Cys | 28.1 | -0.4 | 4 | 3 | 1 | AMR |
| - | p.Tyr199His | 28.1 | -0.2 | 1 | 0 | 1 | SAS |
| rs779790336 | p.Arg204Thr | 28.9 | -0.2 | 1 | 0 | 1 | SAS |
| rs142443432 | p.Asp206Gly | 22.4 | 0.0 | 52 | 23 | 50 | NFE |
| rs754237613 | p.Tyr207Cys | 26.2 | -0.4 | 1 | 0 | 1 | NFE |
| - | p.Val209Ile | 25.6 | 0.0 | 1 | 1 | 1 | NFE |
| rs148771870 | p.Gly211Arg | 23.9 | -0.3 | 243 | 91 | 165 | NFE |
| rs761269389 | p.Asp216Tyr | 27.3 | 0.2 | 1 | 0 | 1 | NFE |
| rs753164828 | p.Asp216Glu | 27.3 | -0.1 | 2 | 1 | 2 | NFE |
| rs372272603 | p.Arg219Cys | 24.9 | -0.2 | 58 | 17 | 52 | NFE |
| rs759590772 | p.Arg219His | 24.9 | 0.0 | 18 | 11 | 17 | SAS |
| rs774621083 | p.Gly220Ser | 32.6 | -0.1 | 3 | 0 | 1 | AFR |
| - | p.Thr229Ile | 37.5 | -0.1 | 1 | 0 | 1 | NFE |
| - | p.His239Gln | 49.7 | 0.0 | 1 | 0 | 1 | NFE |
| - | p.Ala242Val | 50.9 | 0.4 | 3 | 1 | 3 | NFE |
| rs771769548 | p.Tyr252Cys | 47.9 | -0.3 | 2 | 0 | 2 | NFE |
| rs745514718 | p.Ser257Asn | 56.7 | -0.3 | 1 | 0 | 1 | EAS |
| - | p.Ile259Thr | 57.2 | -0.1 | 1 | 0 | 1 | AFR |
| rs200745906 | p.Pro263Ser | 47.8 | 0.1 | 9 | 0 | 9 | NFE |
| - | p.Gly268Cys | 38.9 | -0.7 | 1 | 0 | 1 | ASJ |
| - | p.Asp269Glu | 36.9 | -0.4 | 0 | 0 | 0 |  |
| rs766319182 | p.Met270Val | 37.7 | -0.1 | 4 | 2 | 4 | NFE |
| - | p.Ser280Tyr | 43.8 | 0.0 | 1 | 1 | 1 | NFE |
| rs763264372 | p.Gln287Arg | 49.6 | 0.0 | 1 | 0 | 1 | NFE |
| - | p.Gln287Lys | 49.6 | -0.1 | 1 | 0 | 1 | AMR |
| rs763994205 | p.Asn290His | 39.9 | 0.0 | 2 | 1 | 1 | SAS |
| rs756358940 | p.Ile291Lys | 36.3 | 0.0 | 3 | 2 | 3 | NFE |
| - | p.Asp292Asn | 35.0 | -0.5 | 1 | 0 | 1 | NFE |
| rs776226831 | p.Asp295Gly | 34.8 | -0.1 | 7 | 1 | 6 | NFE |
| rs772143709 | p.Met297Leu | 27.6 | -0.3 | 1 | 0 | 1 | AFR |
| rs745744395 | p.Met297Ile | 27.6 | -0.2 | 1 | 1 | 1 | NFE |
| rs772143709 | p.Met297Val | 27.6 | -0.1 | 0 | 0 | 0 |  |
| rs773936807 | p.Gln300Arg | 29.5 | -0.1 | 2 | 0 | 2 | NFE |
| rs749750821 | p.Asp303Asn | 22.0 | -0.8 | 2 | 0 | 2 | SAS |
| rs766518362 | p.Phe308Leu | 17.0 | -0.5 | 1 | 0 | 1 | NFE |
| rs780574871 | p.Glu312Lys | 12.5 | -0.4 | 3 | 1 | 2 | AFR |
| rs759579097 | p.Gly326Glu | 5.5 | 1.0 | 1 | 0 | 1 | AFR |
| rs143936283 | p.Glu329Gly | 4.1 | -0.4 | 5 | 0 | 4 | NFE |
| rs185525294 | p.Met332Leu | 11.2 | -0.2 | 3 | 2 | 3 | AFR |
| rs762357444 | p.Gly337Arg | 20.2 | 0.0 | 1 | 1 | 1 | NFE |
| - | p.Asn338Ser | 22.9 | 0.0 | 3 | 1 | 2 | AMR |
| rs772710779 | p.Val339Gly | 20.0 | -0.2 | 1 | 1 | 1 | SAS |
| rs138390800 | p.Lys341Arg | 18.1 | 0.0 | 53 | 10 | 46 | AFR |
| - | p.Pro346Ser | 18.9 | 0.0 | 1 | 1 | 1 | NFE |
| rs370610075 | p.Gly352Val | 5.4 | -1.1 | 2 | 0 | 2 | NFE |
| - | p.Asp355Asn | 3.5 | -1.3 | 2 | 0 | 2 | NFE |
| - | p.Met360Leu | 16.0 | -0.1 | 1 | 1 | 1 | NFE |
| rs758568640 | p.Met366Thr | 28.4 | -0.1 | 2 | 0 | 2 | NFE |
| - | p.Asp368Asn | 23.6 | -0.4 | 1 | 0 | 1 | AMR |
| - | p.His374Arg | 20.1 | 0.0 | 1 | 0 | 1 | AMR |
| - | p.Glu375Asp | 16.6 | -0.2 | 3 | 1 | 3 | AMR |
| rs767462182 | p.Gly377Glu | 17.3 | -0.5 | 1 | 1 | 1 | NFE |
| rs142984500 | p.His378Arg | 15.3 | 0.1 | 15 | 4 | 15 | NFE |
| rs751572714 | p.Gln388Leu | 6.6 | -0.1 | 4 | 3 | 4 | AMR |
| rs762890235 | p.Pro389His | 8.1 | -0.1 | 7 | 2 | 4 | AMR |
| - | p.Asn397Asp | 20.9 | -0.1 | 2 | 0 | 2 | NFE |
| rs772619843 | p.Glu398Lys | 22.5 | -0.4 | 2 | 1 | 2 | NFE |
| - | p.Gly405Glu | 22.7 | -0.1 | 1 | 1 | 1 | NFE |
| - | p.Lys419Thr | 33.1 | 0.0 | 1 | 1 | 1 | EAS |
| - | p.Ile421Thr | 26.4 | -0.2 | 1 | 1 | 1 | NFE |
| - | p.Pro426Ala | 38.8 | 0.0 | 1 | 0 | 0 |  |
| - | p.Asp427Tyr | 41.7 | 0.1 | 2 | 1 | 2 | AFR |
| - | p.Gln429Lys | 44.4 | 0.0 | 1 | 1 | 1 | SAS |
| - | p.Asn437His | 42.2 | 0.1 | 0 | 0 | 0 |  |
| rs764772589 | p.Thr445Met | 34.1 | -0.1 | 1 | 1 | 1 | SAS |
| rs776328956 | p.Val447Phe | 38.5 | 0.9 | 10 | 1 | 4 | FIN |
| rs763655186 | p.Gly448Glu | 39.7 | -0.6 | 1 | 1 | 1 | EAS |
| rs760321012 | p.Leu450Val | 35.9 | -0.2 | 1 | 0 | 1 | NFE |
| rs775744448 | p.Met455Ile | 39.9 | -0.1 | 1 | 1 | 1 | SAS |
| - | p.Trp461Arg | 34.1 | -0.3 | 1 | 0 | 1 | AMR |
| rs772445388 | p.Val463Ile | 37.7 | -0.5 | 1 | 1 | 1 | SAS |
| rs774978137 | p.Gly466Trp | 37.7 | 0.0 | 1 | 0 | 1 | EAS |
| - | p.Glu467Lys | 40.4 | -0.1 | 4 | 1 | 1 | AMR |
| rs191860450 | p.Ile468Val | 42.6 | -0.5 | 142 | 40 | 138 | EAS |
| - | p.Lys481Asn | 42.4 | -0.1 | 0 | 0 | 0 |  |
| rs748359955 | p.Arg482Gln | 49.3 | -0.1 | 2 | 1 | 2 | NFE |
| rs779752560 | p.Glu483Asp | 49.2 | 0.1 | 7 | 0 | 7 | AMR |
| rs200973492 | p.Val488Ala | 49.7 | 0.0 | 1 | 0 | 1 | AFR |
| rs750415079 | p.Val491Met | 50.7 | -0.4 | 1 | 1 | 1 | NFE |
| rs765152220 | p.Asp494Val | 49.7 | -0.2 | 8 | 3 | 4 | SAS |
| rs140473595 | p.Ala501Thr | 36.2 | 0.5 | 2 | 0 | 2 | NFE |
| - | p.Phe504Leu | 26.3 | -0.6 | 1 | 0 | 1 | NFE |
| rs760281053 | p.Phe504Ile | 26.3 | -0.3 | 2 | 0 | 2 | NFE |
| - | p.His505Arg | 28.6 | 0.0 | 1 | 0 | 1 | EAS |
| rs775181355 | p.Val506Ala | 33.0 | 0.1 | 1 | 1 | 1 | NFE |
| rs779199005 | p.Tyr510His | 23.2 | -0.1 | 0 | 0 | 0 |  |
| rs868506911 | p.Arg514Gln | 23.4 | -0.1 | 0 | 0 | 0 |  |
| - | p.Arg518Thr | 27.3 | 0.4 | 1 | 1 | 1 | SAS |
| - | p.Thr519Ile | 30.3 | 0.0 | 1 | 1 | 1 | NFE |
| rs759200680 | p.Tyr521His | 26.7 | -0.3 | 1 | 0 | 1 | NFE |
| rs763593286 | p.Ala532Thr | 28.5 | 0.1 | 10 | 5 | 10 | SAS |
| rs773746588 | p.Lys534Arg | 32.6 | -0.1 | 1 | 1 | 1 | NFE |
| - | p.Pro538Leu | 38.7 | 0.1 | 1 | 0 | 1 | SAS |
| - | p.Lys541Ile | 35.8 | -0.1 | 1 | 0 | 1 | AFR |
| rs769821600 | p.Ile544Asn | 25.7 | -0.2 | 1 | 0 | 1 | NFE |
| rs756905974 | p.Asn546Ser | 25.3 | -0.1 | 1 | 0 | 1 | SAS |
| rs761944150 | p.Asn546Asp | 25.3 | 0.0 | 4 | 1 | 3 | AFR |
| rs373025684 | p.Ser547Cys | 24.3 | -0.2 | 43 | 19 | 36 | NFE |
| rs373025684 | p.Ser547Phe | 24.3 | 0.0 | 1 | 0 | 1 | EAS |
| - | p.Arg559Ser | 11.7 | -0.1 | 1 | 0 | 1 | AMR |
| rs375352455 | p.Ser563Leu | 15.2 | 0.1 | 1 | 0 | 1 | NFE |
| rs750955307 | p.Pro565Ser | 21.6 | 0.0 | 1 | 1 | 1 | NFE |
| - | p.Thr567Ala | 25.4 | 0.0 | 1 | 0 | 1 | NFE |
| - | p.Val573Ala | 25.4 | 0.1 | 2 | 1 | 2 | EAS |
| - | p.Val574Ile | 28.4 | -0.2 | 1 | 1 | 1 | SAS |
| - | p.Gly575Arg | 30.3 | -0.1 | 1 | 0 | 1 | NFE |
| rs372924787 | p.Arg582Ser | 38.7 | 0.0 | 1 | 0 | 1 | AFR |
| rs150172355 | p.Arg582Lys | 38.7 | 0.0 | 2 | 0 | 2 | NFE |
| - | p.Leu585Pro | 39.8 | -0.7 | 1 | 0 | 1 | EAS |
| rs768685219 | p.Asn586Tyr | 40.9 | 0.0 | 1 | 0 | 1 | FIN |
| rs760929786 | p.Phe588Ser | 38.6 | -0.4 | 1 | 0 | 1 | AMR |
| - | p.Glu589Gly | 43.9 | -0.1 | 1 | 0 | 1 | FIN |
| rs140857723 | p.Thr593Asn | 49.6 | -0.1 | 1 | 0 | 1 | AFR |
| rs148036434 | p.Leu595Val | 47.7 | -0.2 | 2 | 1 | 2 | NFE |
| rs145437639 | p.Asp597Glu | 54.8 | 0.0 | 13 | 4 | 11 | AFR |
| - | p.Thr608Ile | 53.9 | 0.0 | 1 | 1 | 1 | SAS |
| rs747988885 | p.Asp609Asn | 57.4 | 0.0 | 2 | 1 | 1 | AFR |
| rs201715513 | p.Ala614Ser | 60.5 | 0.0 | 35 | 17 | 7 | FIN |

**Supplementary Table 3.**
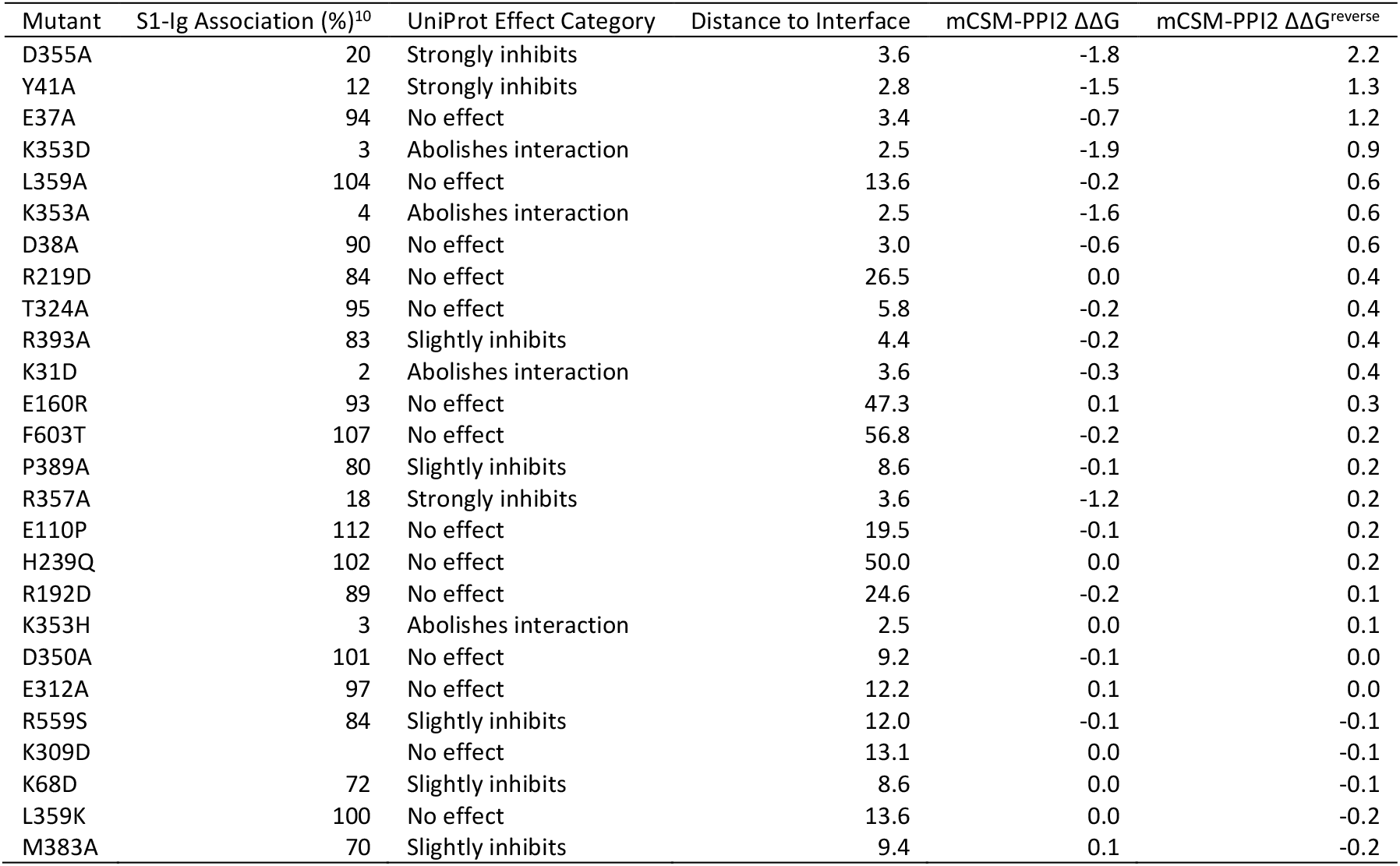
Validation of mCSM-PPI2 with published ACE2 mutant binding assay data.

**Supplementary Figure 1.**
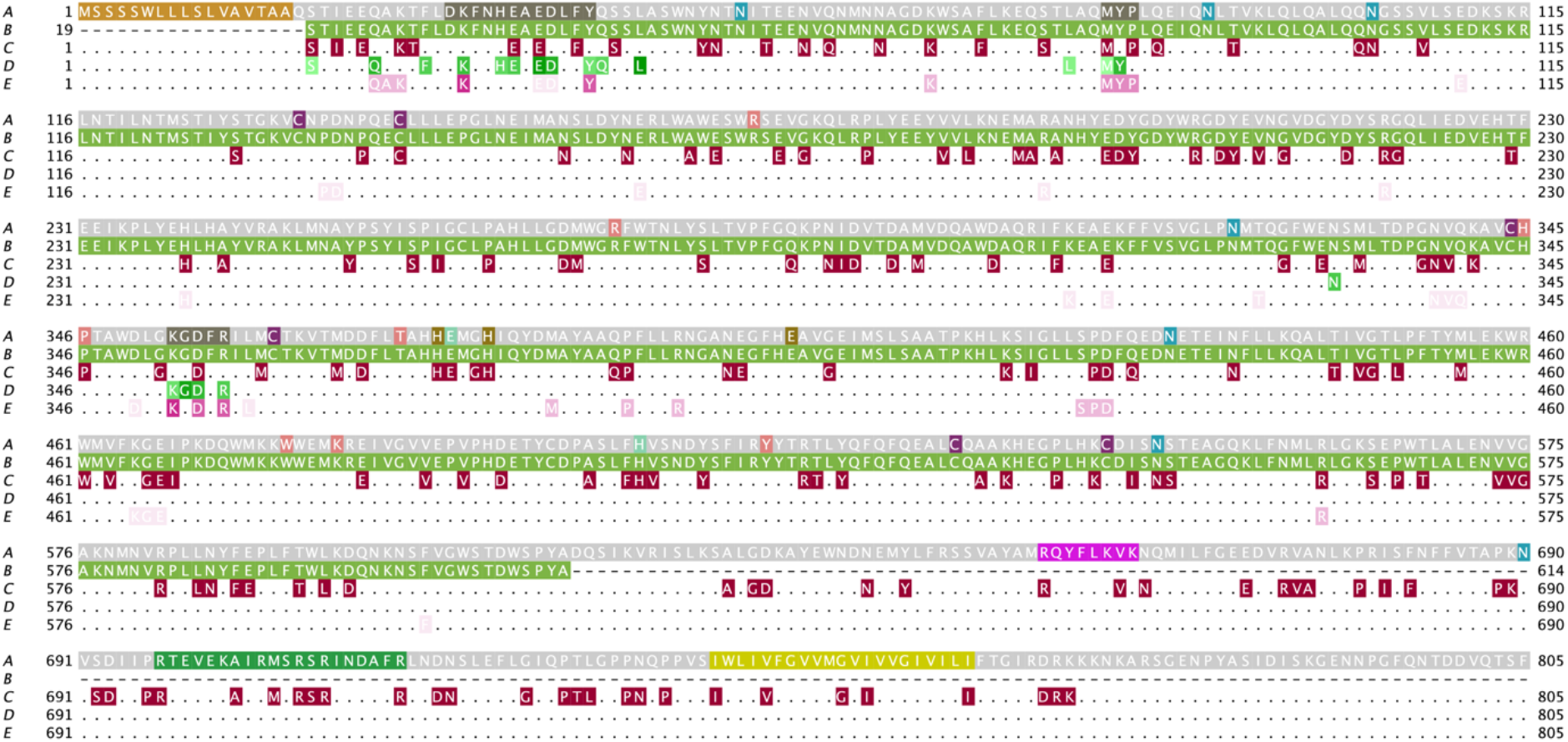
Functional residues in ACE2 and their overlap with missense variants and mutagenesis assays. Each track is the ACE2 sequence with a different set of annotations. A. UniProt annotations. Highlights: signal peptide (orange), SARS-CoV binding (dark grey, positions 30-41, 82-84 and 353-357), glycosylation site (blue) disulphide bond (purple), other binding (pink), metal binding (sand), essential for ADAM17 processing (magenta, 652-659), essential for TMPRSS2 and TMPRSS11D processing (green, 697-716) and transmembrane region (yellow, positions 741-761) B. structural coverage in PDB ID: 6vw1. C. gnomAD missense variants (dark red). D. SARS-CoV-2 S-protein interacting sites (light green to green, proportional to minimum distance to S-protein partner). E. residues with a SARS-CoV interaction mutagenesis assays (light pink to dark pink proportional to effect on ACE2-S affinity). For example, the figure shows that S19 is the first residue resolved in PDB ID: 6vw1 (track B), has a gnomAD missense variant (track C) and is in contact with SARS-CoV-2 S (track D). Data retrieved from UniProt^13^, PDB and UCSF Chimera^16^ from the Jalview^14^ interface. Figure created with Jalview^14^

**Supplementary Figure 2.**
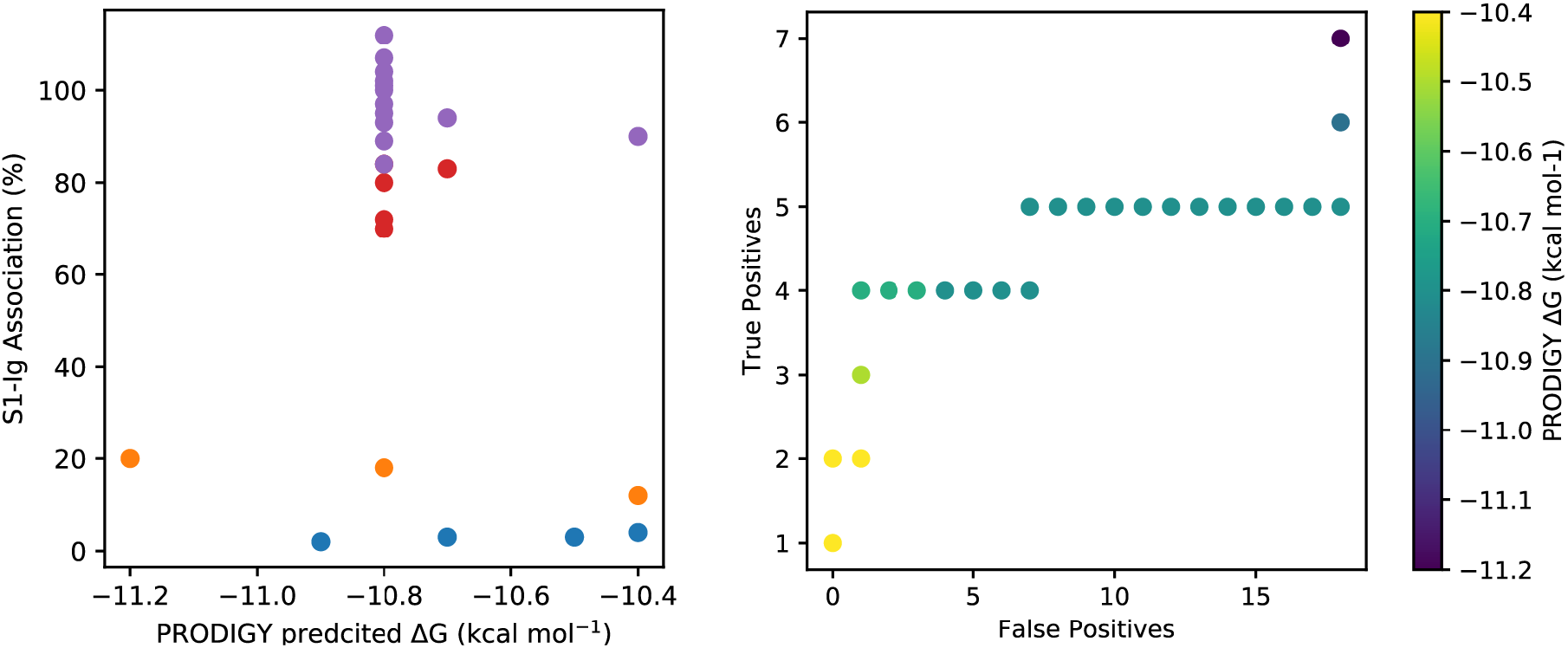
PRODIGY^37^ predictions are inconsistent for ACE2 mutants with SARS-CoV experimental binding assay data^10^. PRODIGY results for mutant models generated by mCSM-PPI2^12^ from PDB ID: 2ajf^8^.

**Supplementary Figure 3.**
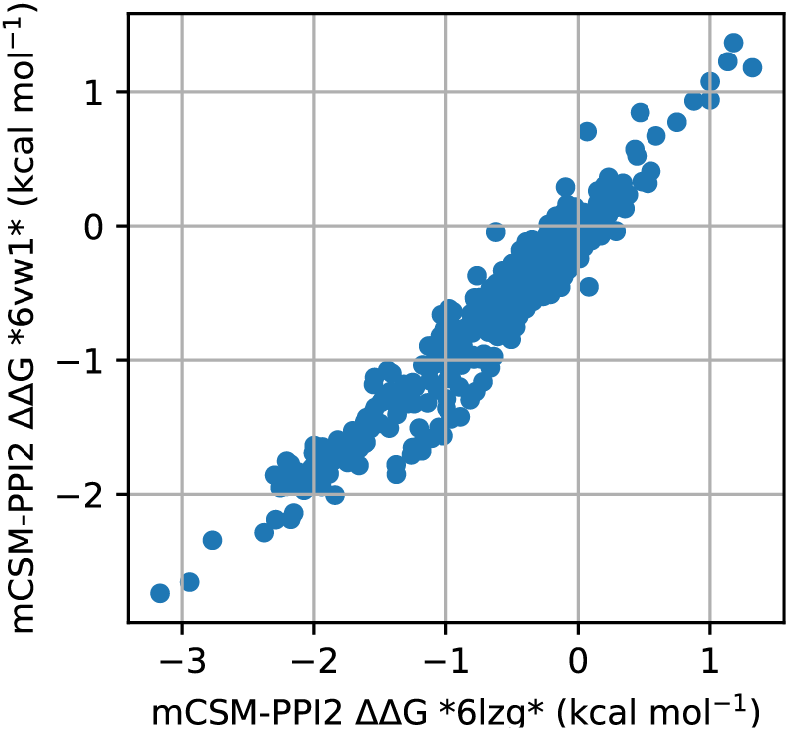
Comparison of mCSM-PPI2^12^ predicted ΔΔG for ACE2 saturation mutagenesis scan (437 mutations at 23 sites) computed for different crystal structures of the ACE2-SARS-CoV-2 S RBD complex, PDB 6vw1^9^ and 6lzg^38^.

**Supplementary Figure 4.**
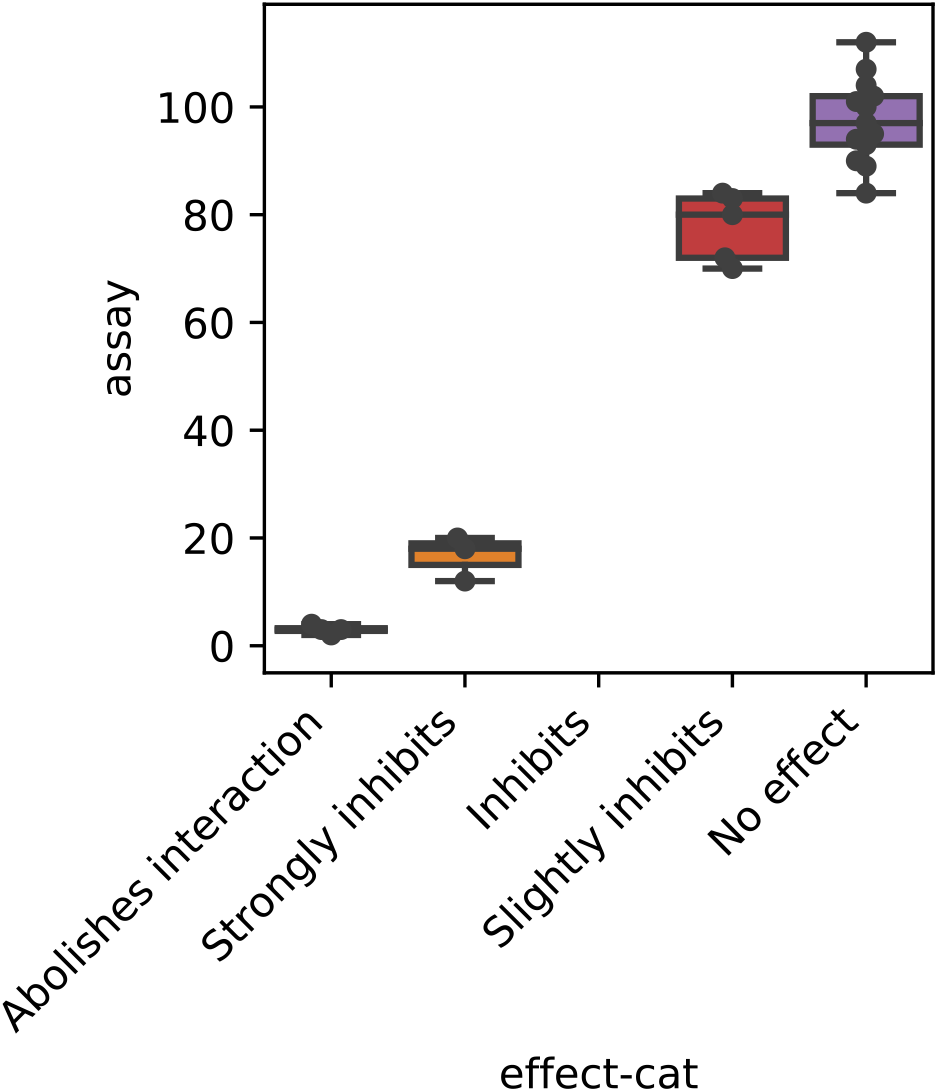
Comparison of mutagenesis assay results from Li et al.^10^ transcribed with WebPlotDigitizer 4.2^40^ with UniProt qualitative annotations.

Supplementary Data File 1. mCSM-PPI saturation mutagenesis data for PDB ID: 6vw1.

